# Elevated phagocytic capacity directs innate spinal cord repair

**DOI:** 10.1101/2024.06.11.598515

**Authors:** Dana Klatt Shaw, Vishnu Muraleedharan Saraswathy, Anthony R. McAdow, Lili Zhou, Dongkook Park, Ridim Mote, Aaron N. Johnson, Mayssa H. Mokalled

## Abstract

Immune cells elicit a continuum of transcriptional and functional states after spinal cord injury (SCI). In mammals, inefficient debris clearance and chronic inflammation impede recovery and overshadow pro-regenerative immune functions. We found that, unlike mammals, zebrafish SCI elicits transient immune activation and efficient debris clearance, without causing chronic inflammation. Single-cell transcriptomics and inducible genetic ablation showed zebrafish macrophages are highly phagocytic and required for regeneration. Cross-species comparisons between zebrafish and mammalian macrophages identified *transcription and immune response regulator* (*tcim*) as a macrophage-enriched zebrafish gene. Genetic deletion of zebrafish *tcim* impairs phagocytosis and regeneration, causes aberrant and chronic immune activation, and can be rescued by transplanting wild-type immune precursors into *tcim* mutants. Conversely, genetic expression of human *TCIM* accelerates debris clearance and regeneration by reprogramming myeloid precursors into activated phagocytes. This study establishes a central requirement for elevated phagocytic capacity to achieve innate spinal cord repair.

## INTRODUCTION

Spinal cord injury (SCI) elicits multicellular and systemic complications that cause permanent neurological deficits in mammals (Matthews et al., 1979; Sofroniew, 2018; Wilkins, 1964). As the first line of defense against neural damage, immune cells are activated as early as 2-3 days and persist years after human SCI (Carlson et al., 1998; Fleming et al., 2006; Means and Anderson, 1983). Immune activation in the central nervous system (CNS) engages tissue-resident microglia and blood-derived leukocytes. Following SCI, lesion-adjacent microglia phagocytose cellular debris and secrete pro-inflammatory cytokines that recruit infiltrating immune cells into the spinal cord (SC) (Hakim et al., 2021; Kigerl et al., 2009). Distinct populations of innate and adaptive immune cells, which do not cross the blood SC barrier under homeostatic conditions, are sequentially recruited to SC lesions (Ankeny et al., 2006; Beck et al., 2010; Popovich et al., 1996; Popovich et al., 1997). Yet, the timeline, composition, and activation profiles of immune cells are infinitely diverse depending on the type, severity, and location of injury (Alexander and Popovich, 2009; Greenhalgh et al., 2018; Hammond et al., 2019; Shechter et al., 2013). Thus, immune cells adopt a range of transcriptional states and functional outputs after mammalian SCI.

Immune cell activation plays dual roles that are challenging to decrypt after mammalian SCI. Consistent with the emergence of an exaggerated inflammatory response that impedes mammalian SC repair, treatment with anti-inflammatory drugs or broad depletion of microglia/macrophages improves functional recovery after mouse SCI (Popovich et al., 1999). Sustained, chronic inflammation is a chief contributor to secondary neurotoxicity, glial cell reactivity, and the recruitment of scar-depositing fibroblasts into the lesion (Dorrier et al., 2021). On the other hand, recently developed tools that enabled targeted microglia/macrophage ablation in acute SCI revealed a requirement for early immune activation in mice (Brennan et al., 2022; Greenhalgh *et al*., 2018; Shechter et al., 2009; Uderhardt et al., 2019). The advent of single-cell transcriptomics provided molecular evidence that microglia/macrophages are highly plastic cells that exist in a spectrum of context-dependent activation states (Hammond *et al*., 2019; Paolicelli et al., 2022). However, correlations between specific immune profiles and their corresponding phenotypic outputs remain to be drawn. The heterogeneity and complexity of immune responses following mammalian SCI present confounding predicaments to deciphering pro-regenerative immune cell states and mechanisms in mammals.

Adult zebrafish present a valuable model to untangle pro-regenerative immune functions in vertebrates. Like mammals, the zebrafish immune system is comprised of evolutionarily conserved innate myeloid and adaptive lymphocyte branches, in addition to CNS-resident microglia (Stachura and Traver, 2011; Xu et al., 2015). Yet, unlike mammals, adult zebrafish are capable of functional recovery after complete SC transection (Becker et al., 1997; Goldshmit et al., 2012; Jensen et al., 2023; Mokalled et al., 2016). Studies of immune cell SCI responses in zebrafish have been limited to distinct immune subsets and/or to larval stages (Cavone et al., 2021; Hui et al., 2017; Tsarouchas et al., 2018). Establishing an immune requirement for larval SC repair, blocking immune activation in 3-day old zebrafish larvae reduces functional and anatomical regeneration (Cavone *et al*., 2021; Tsarouchas *et al*., 2018). However, developmental migration of microglia into the zebrafish CNS extends past 15 days post-fertilization (Xu *et al*., 2015), while mature zebrafish lymphocytes do not emerge until 1 month post-fertilization (Davidson and Zon, 2004; Willett et al., 1997). In adult zebrafish, regulatory T-cells were shown to be required for SC repair (Hui *et al*., 2017). Yet, a comprehensive resource of leukocyte composition and activation profiles after SCI is required to decipher pro-regenerative immune cell states and mechanisms in adult zebrafish.

Limited phagocytic capacity and inefficient debris clearance are associated with mammalian SCI and inhibit SC repair (Becerra et al., 1995). The extracellular milieu of the lesioned cord is flooded with debris-derived, lipid-containing molecules that are potent inhibitors of axon regeneration (Mar et al., 2016). Molecules released by degenerating oligodendrocytes, including neurite outgrowth inhibitors (Nogo) and myelin-associated glycoproteins (MAG), cause growth cone collapse and neurite retraction (McKerracher et al., 1994; Wang et al., 2015a). Inefficient processing of lipid-rich debris transforms phagocytes into lipid-laden foam cells that persist for months to years in human SC lesions (Fleming *et al*., 2006; Pruss et al., 2011; Tamosaityte et al., 2016). Hyper-inflammatory, foamy macrophages lack the ability to migrate or respond to cellular insults, which contributes to chronic inflammation and limits regenerative capacity (Kigerl *et al*., 2009; Kroner et al., 2014; Ryan et al., 2024; Wang et al., 2015b; Zhu et al., 2017). We posit that highly regenerative vertebrates like zebrafish elicit potent phagocytic properties that enable efficient debris clearing after injury. While devising strategies to enhance phagocytic capacity is a prerequisite to improving recovery after SCI, deciphering if and how phagocytic capacity differs between zebrafish and mammals requires cross-species comparisons and mechanistic investigation.

This study examines pro-regenerative immune responses to SCI in adult zebrafish. We report transient and reversible immune cell activation, peaking at 3 days post-injury (dpi) and clearing by 14 dpi. Lesion-adjacent leukocytes are predominantly composed of infiltrating and tissue-resident macrophages. Genetic macrophage depletion during acute and sub-acute SCI impairs functional and anatomical regeneration. Cross-species comparisons between zebrafish and mammalian macrophages identified *transcription and immune response regulator* (*tcim*) as a macrophage-enriched zebrafish gene. Genetic deletion of zebrafish *tcima* and *b* impairs regeneration and can be rescued by transplanting wild-type immune precursors into *tcim* mutants. Validating mass spectrometry data that implicated Tcim in lipid processing and phagocytosis, *tcima/b* mutants showed excessive accumulation of lipid droplets and reduced phagocytic capacity. Conversely, genetic overexpression of human *TCIM* accelerates debris clearance, lipid processing, and regeneration by reprogramming myeloid precursors into activated phagocytes. This study elucidates fundamental differences in the cellular and molecular profiles of immune cells between regenerative and non-regenerative vertebrates in response to SCI and establishes a requirement for elevated phagocytic capacity to achieve innate SC repair.

## RESULTS

### An acute, transient immune response is activated after SCI in adult zebrafish

The zebrafish immune system is comprised of an innate myeloid branch (neutrophils and macrophages) and an adaptive lymphoid branch (B-and T-cells) that are conserved across vertebrates (Davidson and Zon, 2004; Stachura and Traver, 2011). In addition to myeloid-derived macrophages, the CNS is privileged with tissue-resident microglia that elicit a phagocytic response after CNS injury or disease (Krasemann et al., 2017; Nguyen-Chi et al., 2015; Villani et al., 2019; Wittamer et al., 2011). To assess the identities of the immune cells responding to SCI, we performed complete SC transections on *Tg(lyz:EGFP)* reporter fish that label neutrophils (Hall et al., 2007) or *Tg(lck:lck-EGFP)* fish for T-cell labeling (Langenau et al., 2004), and co-stained with the panleukocyte marker Lcp1 (Redd et al., 2006). Consistent with prior findings in larval zebrafish and mice (Carlson *et al*., 1998; Means and Anderson, 1983; Tsarouchas *et al*., 2018), neutrophils were the first leukocytes to respond to SCI, peaking at 12 hours post-injury (Fig. S1A-B), while T-cells showed maximum infiltration at 14 dpi (Fig. S1C-D). However, only 10.6% of leukocytes were neutrophils at 1 dpi and 14.5% of leukocytes were T-cells at 14 dpi (Fig. S1E), indicating that neutrophils and T-cells comprise small fractions of Lcp1^+^ leukocytes after zebrafish SCI.

We next analyzed Lcp1-stained SC sections from *Tg(mpeg1.1:YFP)* reporter fish to assess the numbers of infiltrating and tissue-resident macrophages between 1 and 56 dpi (Ellett et al., 2011). Prior to injury, the number of *mpeg1.1*^+^ leukocytes, which are presumed to represent microglia in the presence of an intact blood-spinal cord barrier, averaged 22.3 ± 3.3 per section (Fig. 1A-B). Following SCI and barrier disruption, tissue-resident and infiltrating macrophages showed a 12-fold increase from 0 to 3 dpi (265.5 ± 52.4 cells per section) (Fig. 1A-B). Macrophage numbers returned to pre-injury levels at 28 dpi (p=0.22 compared to uninjured) (Fig. 1A-B). *mpeg1.1*^-^Lcp1^+^ leukocytes were present in smaller numbers at the lesion site and rarely infiltrated beyond 300 µm into the rostral or caudal SC parenchyma (Fig. 1B). Confirming these results, sorting *mpeg1.1*^+^ cells from *Tg(mpeg1.1:YFP)* SCs isolated 3.73-fold more cells at 5 dpi compared to uninjured controls (Fig. S1F). Thus, while elevated leukocytes are known to persist into the chronic injury phase in mouse and human SC tissues after SCI (Gadani et al., 2015; Gillespie and Ruitenberg, 2022), immune responses to SCI are transient in zebrafish. These studies indicated that infiltrating and tissue-resident macrophages are the predominant leukocyte populations that respond to SCI in adult zebrafish and suggested that acute phagocyte activation is an important feature of SC regeneration.

**Figure 1.**
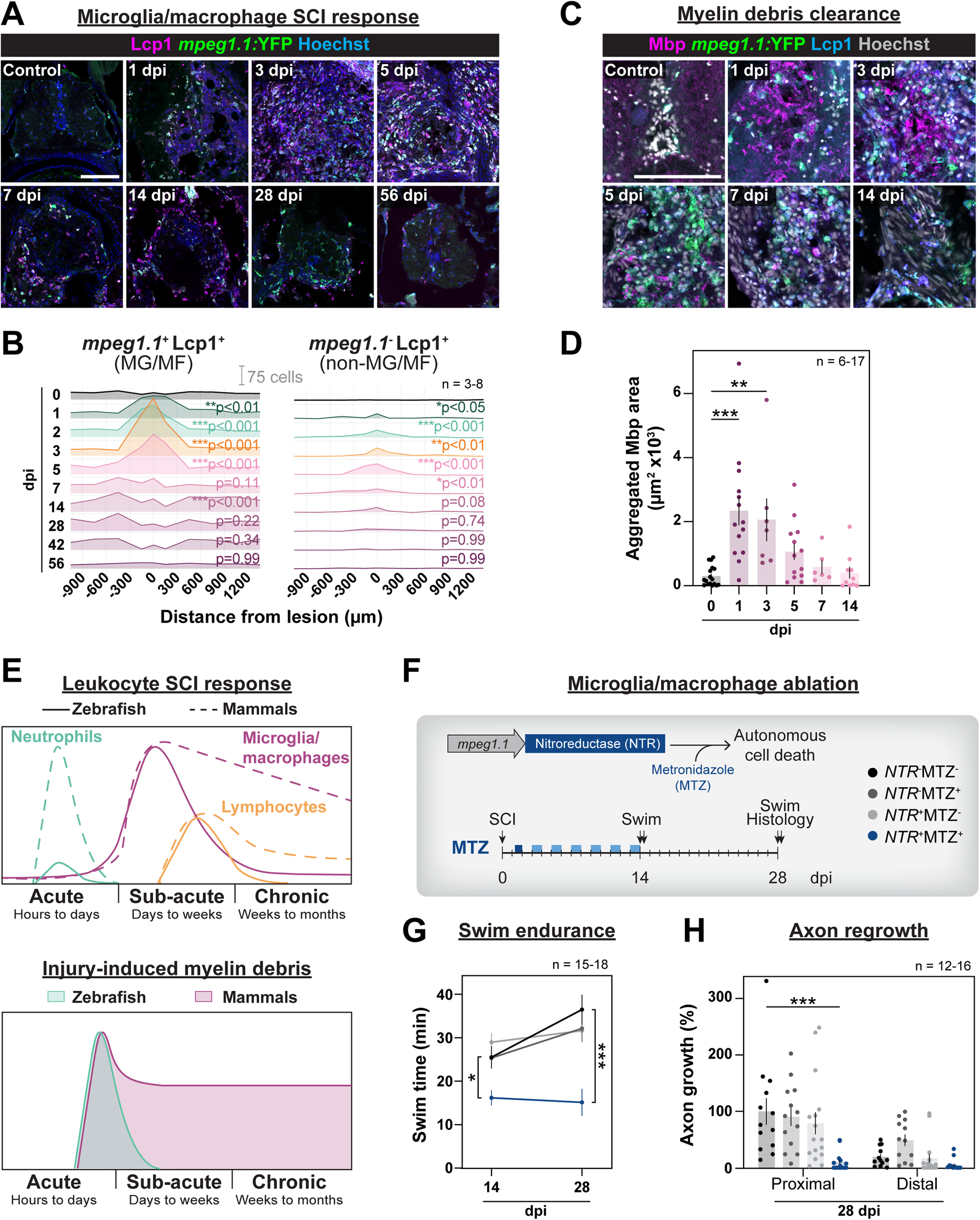
A transient microglia/macrophage response is required for SC regeneration. **(A)** Immunostaining for the pan-leukocyte marker L-plastin (Lcp1) in *Tg(mpeg1.1:YFP)* zebrafish. *Tg(mpeg1.1:YFP)* is expressed by microglia and macrophages (MG/MF). Cross sections from uninjured SCs (controls) or between 1 and 56 dpi are shown. Scale bar, 100 µm. **(B)** Ridgeline plots of microglia/macrophages (MG/MF, *mpeg1.1:*YFP^+^Lcp1^+^) or non-MG/MF (*mpeg1.1:*YFP^-^ Lcp1^+^) before and after SCI. Scale bar, 75 cells. Distance from the lesion is indicated in µm where negative values indicate rostral levels and positive values indicate caudal levels. Sample sizes: Uninjured (4), 1 dpi (4), 2 dpi (7), 3 dpi (4), 5 dpi (8), 7 dpi (5), 14 dpi (3), 28 dpi (5), 42 dpi (4), 56 dpi (4). Two-way ANOVA (Dunnett’s): *p<0.05, **p<0.01, ***p<0.001. Comparisons are to uninjured controls. **(C,D)** Mbp and Lcp1 staining in *Tg(mpeg1.1:YFP)* zebrafish. Cross sections from uninjured SCs (controls) or between 1 and 14 dpi are shown. Mbp aggregates were determined using a threshold that labeled <1% of the total area in uninjured SC sections. Scale bar, 100 µm. Sample sizes: Uninjured (17), 1 dpi (14), 3 dpi (7), 5 dpi (13), 7 dpi (6), 14 dpi (10). One-way ANOVA (Dunnett’s): **p<0.01, ***p<0.0001. **(E)** Schematic comparison of zebrafish immune responses after SCI (A-B) with previously reported human and mouse immune responses (top), or the prevalence of myelin debris in adult zebrafish (C-D) an in human SCI (bottom). Mammalian data is adapted from (Gadani *et al*., 2015; Gillespie and Ruitenberg, 2022) and (Becerra *et al*., 1995). **(F)** *Tg(mpeg1.1:NTR-IRES-EGFP)* fish were treated with Metronidazole (MTZ) to ablate *mpeg1.1*^+^ cells. 5 mM MTZ followed by 1 mM MTZ pulses were used to maintain depletion between 0 and 14 dpi. SC regeneration was assessed by functional recovery swims and histology for anatomical regeneration at 14 and 28 dpi. **(G)** Swim endurance of microglia/macrophage-depleted fish was measured at 14 and 28 dpi. Sample sizes: NTR^-^MTZ^-^ (16), NTR^+^MTZ^-^ (15), NTR^-^MTZ^+^ (18), NTR^+^MTZ^+^ (16). Two-way ANOVA (Dunnett’s): *p<0.05, ***p<0.001. All samples were compared to time-matched NTR^-^MTZ^-^. **(H)** Anatomical regeneration of microglia/macrophage depleted fish was measured by anterograde axon labeling at 28 dpi. Sample sizes: NTR^-^MTZ^-^ (13), NTR^+^MTZ^-^ (16), NTR^-^MTZ^+^ (13), NTR^+^MTZ^+^ (15). Two-way ANOVA (Dunnett’s): ***p<0.001. Samples were compared to NTR^-^MTZ^-^.

### Myelin debris is efficiently cleared after zebrafish SCI

To assess the extent of debris clearance following zebrafish SCI, we performed Myelin basic protein (Mbp) and Lcp1 staining on SC sections from *Tg(mpeg1.1:YFP)* fish. Mbp, which surrounded myelinated axons in uninjured SC sections, formed debris aggregates within the lesion site after SCI (Fig. 1C). By thresholding dense Mbp^+^ punctae (Fig. S1G), we found minimal Mbp^+^ aggregation in uninjured SC sections (0.29% of the total imaged area). Following SCI, Mbp^+^ aggregation increased 8.1-(p<0.0001) and 6.9-fold (p=0.0031) at 1 and 3 dpi, respectively (Fig. 1C-D). By 5 dpi, Mbp^+^ aggregation was statistically comparable to controls (p=0.24) (Fig. 1C-D). Thus, compared to the immune and phagocytic responses after mammalian SCI (Becerra *et al*., 1995; Gadani *et al*., 2015; Gillespie and Ruitenberg, 2022), transient leukocyte activation and efficient myelin debris clearance are unique features that correlate with elevated regenerative capacity after zebrafish SCI (Fig. 1E).

### *mpeg1.1*^+^ phagocytes are required for SC regeneration

To examine whether transient phagocyte activation is required for SC regeneration, we depleted *mpeg1.1*^+^ phagocytes using the *Tg(mpeg1.1:NTR-IRES-EGFP)* transgene (Hughes and Appel, 2020). With this inducible genetic ablation system, *mpeg1.1*-driven expression of the Nitroreductase (NTR) enzyme catalyzes the reduction of the Metronidazole (MTZ) prodrug into a cytotoxic product, resulting in cell-autonomous ablation of *mpeg1.1*^+^ cells (Curado et al., 2008). Because macrophages undergo rapid replacement after depletion, continuous MTZ pulses were required to sustain *mpeg1.1*^+^ cell depletion. *Tg(mpeg1.1:NTR-IRES-EGFP)* zebrafish were treated with 5 mM MTZ for 18 hours at 1 dpi, followed by 1 mM treatments for 24 hours every other day starting at 3 dpi (Fig. 1F). This pulsed MTZ treatment decreased Lcp1^+^ cells by 74.4%, 73.1%, and 48.5% at 3, 5, and 7 dpi, respectively (Fig. S1H). To determine if acute to sub-acute activation of *mpeg1.1*^+^ phagocytes is required for SC regeneration, we performed SC transections on *Tg(mpeg1.1:NTR-IRES-EGFP)* fish, pulsed MTZ from 1 to 14 dpi, and assessed functional and anatomical regeneration. Macrophage-depleted animals, which exhibited a 36.5% (p=0.0428) decrease in swim endurance at 14 dpi compared to vehicle-treated transgene-negative controls, failed to recover between 14 and 28 dpi (p<0.0001) (Fig. 1G). Using anterograde axon tracing and Gfap immunostaining, proximal axon regrowth and glial bridging across the lesion were significantly reduced in macrophage-ablated animals compared to control cohorts at 28 dpi (Fig. 1H and S1I-K) (Mokalled *et al*., 2016). Thus, although MTZ removal at 14 dpi was sufficient for *mpeg1.1*^+^ cells to repopulate SC tissues, delayed macrophage emergence was not sufficient to direct regeneration between 14 and 28 dpi (Fig. 1G-H and S1I-K). These results indicated *mpeg1.1*^+^ phagocytes are acutely required for SC regeneration in adult zebrafish.

### Comparative transcriptomics identifies a unique transcriptional signature in zebrafish phagocytes

Concomitant with the emergence and requirement for *mpeg1.1*^+^ phagocytes, cellular debris are efficiently cleared following zebrafish SCI (Fig. 1). We postulated that zebrafish macrophages elicit a unique transcriptional signature that directs elevated phagocytic capacity and pro-regenerative functions. To test this hypothesis, we analyzed the profiles of zebrafish and mouse immune cells after SCI using previously generated single-cell RNA-sequencing datasets (Fig. 2A) (Milich et al., 2021; Saraswathy et al., 2023). Two zebrafish leukocyte clusters, referred to as M/M1 and M/M2, expressed canonical microglia/macrophage markers, including *mpeg1.1, csf1ra, csf1rb,* and *lgals3bpb* (Fig. 2A and S2A). M/M1 represented 99.5% and 94.6% of all zebrafish microglia/macrophages in uninjured samples and at 7 dpi, respectively. By filtering top M/M1 markers for genes that are upregulated at 7 dpi compared to uninjured SCs, we identified 331 macrophage-enriched and injury-induced zebrafish genes (Fig. 2A and Table S1). Out of the 331 M/M1-enriched zebrafish genes, 134 genes were downregulated in mouse microglia at the same time point (Fig. 2A). Intriguingly, the injury-induced gene signature widely differs between zebrafish and mice, with only 15 genes upregulated in myeloid phagocytes at 7 dpi in both species (Fig. 2A). Gene ontology (GO) analysis showed zebrafish M/M cells enriched for immune-related genes, with “cell activation” being the most statistically significant GO term (-log_10_(p-value)=11.1) (Fig. 2A and Table S2). The 27 genes implicated in immune cell activation included immune-related genes, such as *mef2cb* (Wang et al., 2016), *klf6a* (Kramer et al., 2021), and *il4r.1* (Pan et al., 2022). We also identified genes that were not previously implicated in regeneration or the CNS including *erythroblast transformation-specific transcription factor 1* (*elf1*), *phospholipase C like 2* (*plcl2*), and *transcription and immune response regulator a* (*tcima*). These comparative studies identified genes that are preferentially enriched in zebrafish microglia/macrophages and likely to be associated with increased phagocytic and regenerative capacities in zebrafish.

**Figure 2.**
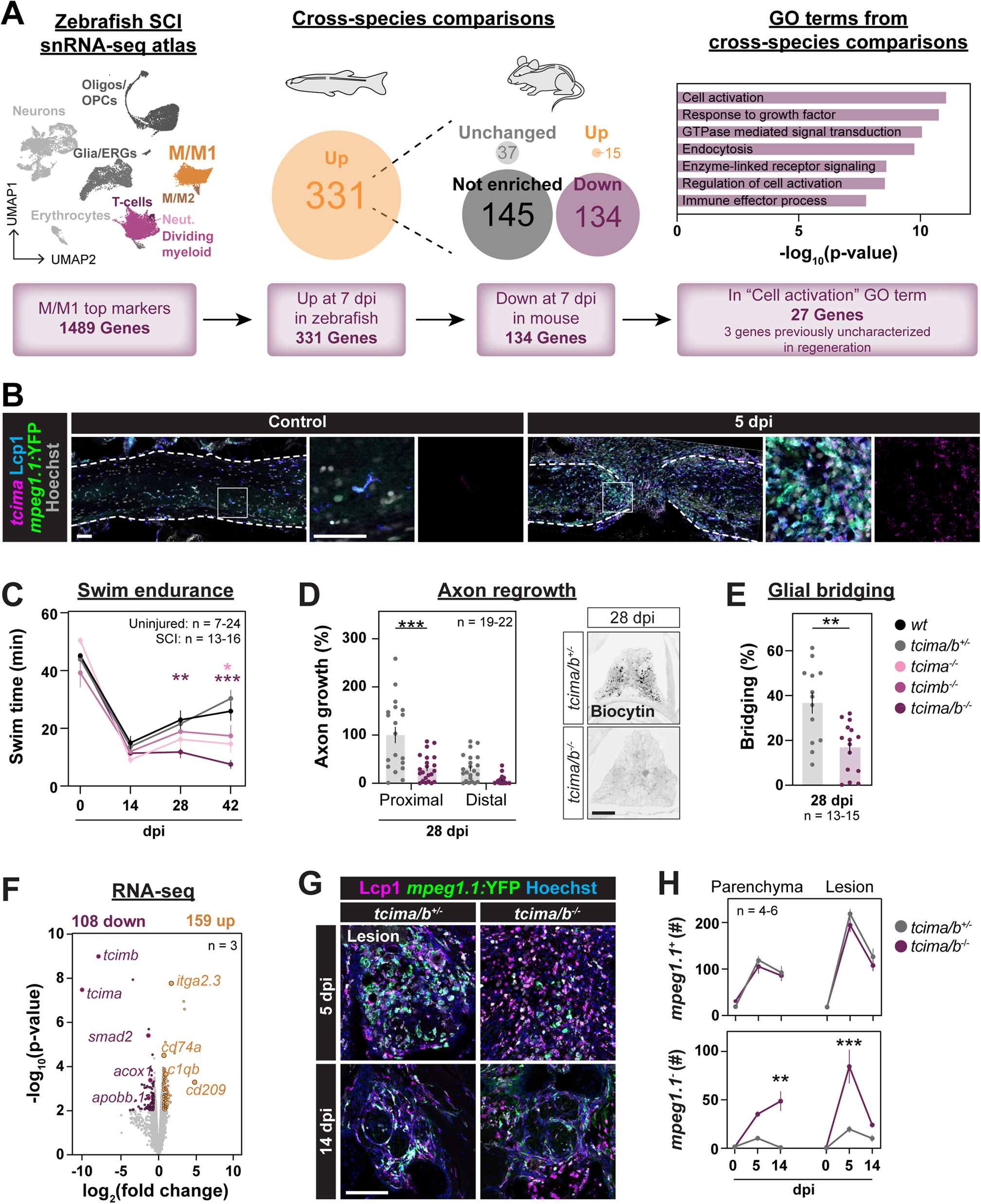
*tcim* is enriched in microglia/macrophages and required for SC regeneration. **(A)** A pipeline to identify zebrafish-enriched immune genes after SCI. An single nuclear RNA-sequencing (snRNA-seq) atlas of SC regeneration in adult zebrafish was recently generated (Saraswathy *et al*., 2023), and mouse microglia data was analyzed (Milich *et al*., 2021). The microglia/macrophage cluster 1 (M/M1) expressed the canonical marker genes *mpeg1.1, csf1ra, csf1rb,* and *lgals3bpb*. M/M1 top marker genes at 7 dpi were filtered for genes that were downregulated at the same time point in mouse microglia. Among 134 genes upregulated at 7 dpi in zebrafish but downregulated in mouse at 7 dpi, 27 genes were related to the GO term “cell activation”. Genes expressed in >10% of cells, enriched with a log_2_(fold enrichment) >0.25 compared to all non-microglia/macrophage cells, and with p-value of <0.01 with the Wilcoxon Rank Sum test were considered enriched. **(B)** HCR *in situ* hybridization of *tcima* and Lcp1 staining were performed on *Tg(mpeg1.1:YFP)* fish. Sagittal SC sections from uninjured or 5 dpi animals were used. Scale bars, 50 µm. **(C)** Swim endurance following tcim loss-of-function. Wild-type (*wt*), *tcima/b*^+/-^, *tcima*^-/-^, *tcimb*^-/-^, and *tcima/b*^-/-^ animals were used. Swim assays were performed pre-injury (0 dpi) or at 14, 28, and 42 dpi. Sample sizes (pre-injury): wild-type (9), *tcima/b*^+/-^ (24), *tcima*^-/-^ (21), *tcimb*^-/-^ (7), and *tcima/b*^-/-^ (17). Sample sizes (SCI): wild-type (15), *tcima/b*^+/-^ (13), *tcima*^-/-^ (16), *tcimb*^-/-^ (14), and *tcima/b*^-/-^ (16). One-way ANOVA (Sidak’s): *p<0.05, **p<0.01, ***p<0.001. Comparisons are to timepoint-matched wild-type siblings. **(D,E)** Anatomical regeneration of *tcima/b^-/-^* and control fish was measured by anterograde axon labeling (D) or glial bridging quantification (E). Representative images of biocytin-traced axons are shown in D. Scale bar in D, 100 µm. Sample sizes (axon regrowth): *tcima/b*^+/-^ (19) and *tcima/b*^-/-^ (22). One-way ANOVA (Sidak’s): ***p<0.001. Sample sizes (glial bridging): *tcima/b*^+/-^ (13) and *tcima/b*^-/-^ (15). Unpaired t- test: **p<0.01. **(F)** Bulk RNA-seq of *tcima/b*^-/-^ and control SCs at 5 dpi. Sample sizes represent 9-10 SCs per pool and 3 triplicate pools were used. **(G,H)** Lcp1 staining on *Tg(mpeg1.1:YFP); tcima/b^-/-^*and *Tg(mpeg1.1:YFP); tcima/b^+/-^ fish.* Cross sections at the lesion core are shown at 5 and 14 dpi. Scale bar, 100 µm. Parenchyma indicates 150 µm caudal to the lesion core. Sample sizes: *tcima/b*^+/-^ uninjured (6), *tcima/b*^-/-^ uninjured (4), *tcima/b*^+/-^ 5 dpi (6), *tcima/b*^-/-^ 5 dpi (6), *tcima/b*^+/-^ 14 dpi (5), *tcima/b*^-/-^ 14 dpi (6). Two-way ANOVA (Sidak’s): **p<0.01, ***p<0.001. Comparisons are to time-matched *Tg(mpeg1.1:YFP); tcima/b^+/-^* siblings.

### *tcim* expression is upregulated in zebrafish macrophages after SCI

The zebrafish genome harbors the gene paralogs *tcima* and *tcimb*. *tcim* genes encode 101 amino acid-containing peptides, which were shown to regulate cell proliferation and inflammation in cultured HEK and liver cancer stem cells and during zebrafish embryonic development (Kim et al., 2006a; Kim et al., 2009; Kim et al., 2006b; Park et al., 2007; Su et al., 2013; Sunde et al., 2004; Zhu et al., 2015). However, the *in vivo* functions of Tcim during regeneration or within the CNS are unknown. At 7 dpi, zebrafish *tcima* expression is increased in microglia/macrophages (log_2_(fold change)=0.64), while murine *tcim* is downregulated in microglia (log_2_(fold change)=-0.81) (Fig. S2B) (Milich *et al*., 2021; Saraswathy *et al*., 2023). To confirm these observations, we performed complete SC transections on *Tg(mpeg1.1:YFP)* reporter zebrafish, stained for Lcp1, and performed HCR *in situ* hybridization for *tcima* at 5 dpi and in uninjured animals. Concomitant with the emergence of microglia/macrophages in the lesion site and surrounding parenchyma, *tcima* showed minimal baseline expression in uninjured SC tissues but was upregulated in *mpeg1.1^+^*Lcp1^+^ cells at 5 dpi (Fig. 2B and S2C). The *tcima* paralog, *tcimb,* was not identified as a unique macrophage marker. However, as zebrafish paralogs have been shown to play redundant functions upon genetic perturbation (El-Brolosy et al., 2019; Klatt Shaw and Mokalled, 2021; Klatt Shaw et al., 2021), we also examined the expression of *tcimb* after SCI. *tcimb* transcripts were expressed in a subset of *mpeg1.1*^+^Lcp1^+^ cells and in *mpeg1.1*^-^Lcp1^-^ cells after SCI (Fig. S2D). These studies showed that zebrafish *tcima* expression is enriched in microglia/macrophages after SCI.

### *tcim* loss-of-function impairs SC regeneration

To determine the role of *tcim* during SC regeneration, we generated CRISPR/Cas9-mediated gene deletions of zebrafish *tcima* and *tcimb* (Fig. S2E-F). To account for putatively redundant or compensatory *tcim* functions, we examined SC regeneration in *tcima*^-/-^;*tcimb*^-/-^ zebrafish (referred to hereafter as *tcima/b* mutants or *tcima*/*b*^-/-^ fish). Single and double *tcima/b* mutants are adult viable and show normal swim endurance prior to injury (Fig. 2C). However, after SCI, swim endurance was reduced by 48.6% (p=0.0087) and 58.7% (p=0.0003) in *tcima/b* mutants compared to wild-type siblings at 28 and 42 dpi, respectively (Fig. 2C). *tcima/b* mutants did not recover swim endurance capacity between 14 and 42 dpi. At 42 dpi, *tcima* loss-of-function elicited an intermediate phenotype compared to *tcima/b* mutants, suggesting a compensatory role for *tcimb* in *tcima* mutants (Fig. 2C). We therefore used *tcima/b* mutants for the remaining loss-of-function studies. As *tcima/b* heterozygous and wild-type siblings showed comparable swim endurance (Fig. 2C), *tcima/b* heterozygous animals were used as controls. By anterograde axon tracing, *tcima/b* mutants exhibited 69.0% (p<0.0001) and 80.5% (p=0.09) reduction in axon regrowth at 28 dpi proximal and distal to the lesion, respectively (Fig. 2D). By Gfap staining, glial bridging decreased by 53.9% (p=0.0011) in *tcima/b* mutants relative to controls (Fig. 2E and S2G). These studies indicated *tcima/b* are required for functional recovery and anatomical regeneration in adult zebrafish.

### *tcim* loss-of-function impairs immune cell clearance following SCI

To survey the molecular mechanisms that underlie impaired SC regeneration in *tcima/b* mutants, we performed SC transections on *tcima/b* mutant and control siblings and collected 4 mm SC tissue sections spanning the lesion site at 5 dpi for bulk RNA-sequencing. We found 159 genes upregulated and 108 genes downregulated in *tcima/b* mutants compared to controls (Fig. 2F and Table S3). By GO analysis, genes upregulated in *tcima/b* mutants were associated with blood development, amino acid metabolism, and immune activation and included pro-inflammatory genes such as *itga2.3*, *cd74a*, *c1qb,* and *cd209* (Fig. 2F, S2H and Table S4) (Bohlson et al., 2014; Peters et al., 2012; Su et al., 2017; Zhou et al., 2006). Conversely, genes downregulated in *tcima/b* mutants were enriched for terms related to cell proliferation, immune homeostasis, and lipid metabolism (Fig. S2I and Table S5). Specifically, *smad2*, an important regulator of immune suppression (Takimoto et al., 2010), and the lipid processing genes *apobb.1* and *acox1* (Chung et al., 2020; Templehof et al., 2021) were markedly downregulated in *tcima/b* mutants (Fig. 2F). Thus, a pro-inflammatory signature is activated in *tcima/b* mutants, suggesting that *tcim* regulates immune cell activation and function following SCI.

To determine immune cell composition upon *tcima/b* loss-of-function, we performed histology for microglia/macrophages and pan-leukocytes in *tcima/b* mutants. *Tg(mpeg1.1:YFP)* fish were crossed into the *tcima/b* mutant background and subjected to SCI and tissue collection at 5 and 14 dpi. The numbers of lesion-associated or parenchymal *mpeg1.1*^+^Lcp1^+^ leukocytes were not changed in *tcima/b* mutants compared to controls at either 5 or 14 dpi (Fig. 2G-H). However, at 5 dpi, the numbers of lesion-associated *mpeg1.1*^-^Lcp1^+^ leukocytes increased from 19.5 ± 3.4 cells in controls to 84.3 ± 17.4 cells per section in *tcima/b* mutants (p<0.0001) (Fig. 2G-H). At 14 dpi and concomitant with the clearance of *mpeg1.1*^-^Lcp1^+^ leukocytes from the lesion site, the numbers of parenchymal *mpeg1.1*^-^Lcp1^+^ leukocytes increased from 0.6 ± 0.4 cells in control fish to 48.7 ± 10.1 cells per section in *tcima/b* mutants (p=0.0042) (Fig. 2G-H). These results underscored aberrant immune cell composition and chronic immune activation in *tcima/b* mutants after SCI.

### Hematopoietic stem cell transplantation rescues impaired regeneration in *tcim* mutants

Global deletion of *tcima/b* impairs SC regeneration and causes excessive immune activation. To determine the cell-specific requirement for *tcima/b*, we combined hematopoietic transplantation protocols (Traver et al., 2004) with microglia/macrophage ablation (Shechter *et al*., 2009; Xu et al., 2020), which allowed us to differentially control the genotypes of immune cells in *tcima/b* mutants. In a first step to deplete microglia/macrophages, we treated *Tg(mpeg1.1:NTR-IRES-EGFP)*; *tcima/b^-/-^*zebrafish with MTZ for 24 hours starting 72 hours prior to transplantation. We then subjected MTZ-treated animals to a split dose of 8Gy/8Gy irradiation 48 and 24 hours before transplantation. This protocol generated microglia and immune-deficient *tcima/b* mutants. Unlike mammalian bone marrow, the zebrafish hematopoietic niche is located in the kidney (Davidson and Zon, 2004). Thus, for hematopoietic transplantation, fluorescently-labeled kidney cells were dissociated for intraperitoneal injection into *tcima/b* mutants (Fig. 3A). Compound *Tg(actb2:membrane-Citrine); tcima/*b*^+/+^* and *Tg(actb2:membrane-Citrine); tcima/*b*^-/-^* donors were used. To assess transplantation efficiency, we quantified the proportions of Citrine^+^Lcp1^+^ cells relative to total Lcp1^+^ leukocytes in uninjured hosts. At 14 days post-transplantation (dpt), wild-type and *tcima/b^-/-^* donor cells accounted for >35% (p<0.0001) and >40% (p<0.0001) of Lcp1^+^ cells in the kidney and SC tissues, respectively (Fig. 3B-C). These results indicated successful transplantation of wild-type and *tcima/b^-/-^* cells and allowed us to selectively modify immune cell genotypes during SC regeneration.

**Figure 3.**
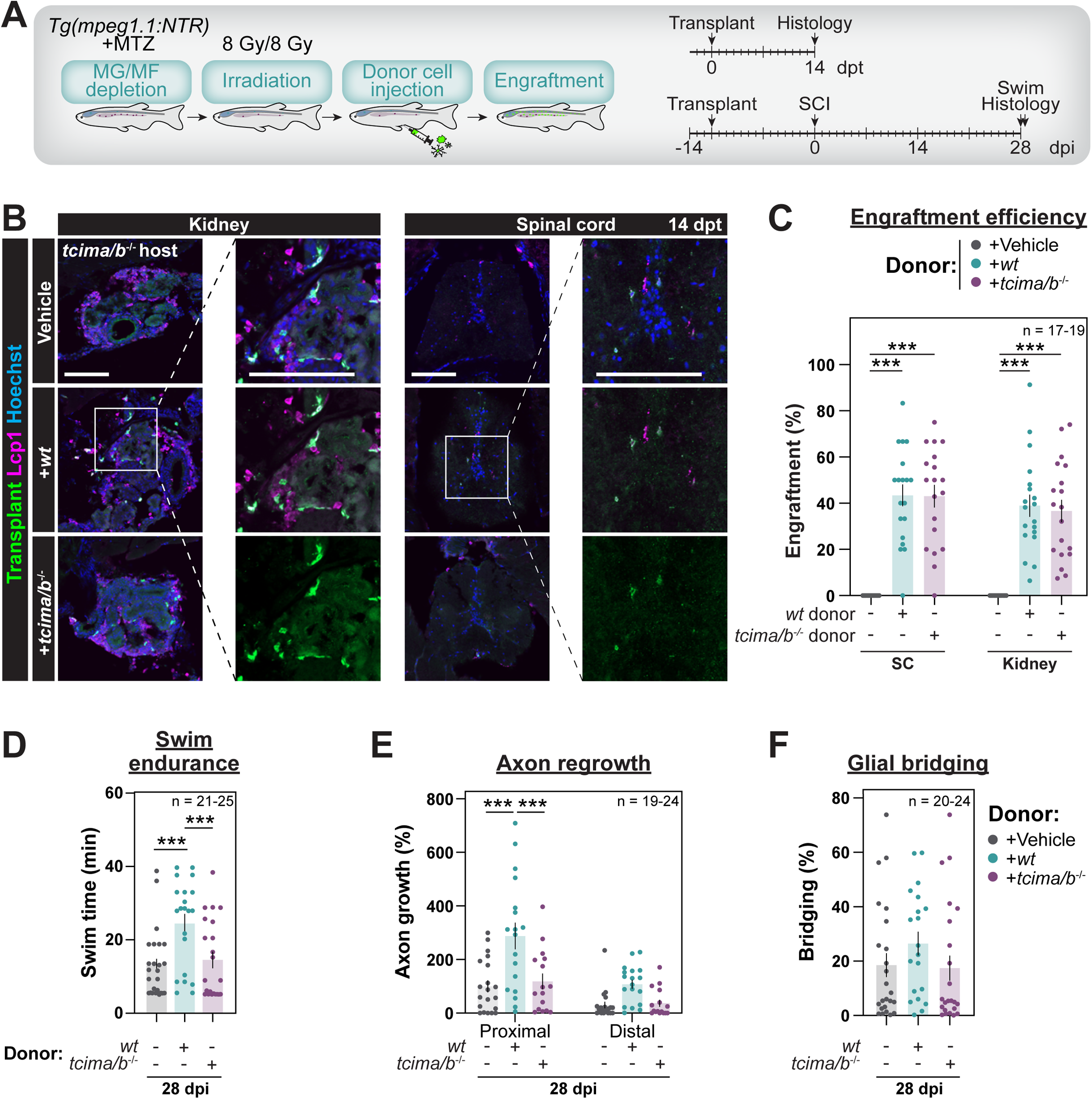
Wild-type hematopoietic cell transplantation rescues SC regeneration in *tcima/b* mutants. **(A)** Experimental timeline to generate immune chimeras via whole kidney marrow replacement. *Tg(mpeg1.1:NTR-IRES-EGFP)* fish were treated for 24 hours with high dose (10 mM) MTZ to deplete microglia/macrophages followed by two split doses of 8 Gy irradiation to deplete the hematopoietic kidney marrow niche. Ubiquitously-labeled whole kidney marrow donor cells were dissociated from *Tg(actb2:membrane-Citrine)* fish. Donor cells were intraperitoneally injected into irradiated hosts for engraftment. **(B,C)** GFP and Lcp1 staining following hematopoietic transplantation. Transplanted *Tg(actb2:membrane-Citrine)* donor cells were detected in the uninjured SC and kidney tissues. Scale bars, 100 µm. Sample sizes: Vehicle (17), wild-type donor (19), *tcima/b*^-/-^ donor (19). One-way ANOVA (Sidak’s): ***p<0.0001. Comparisons are to vehicle-injected controls. **(D)** Swim endurance in transplanted *tcima/b*^-/-^ mutant hosts. Sample sizes: Vehicle (25), wild-type donor (21), *tcima/b*^-/-^ donor (22). One-way ANOVA (Sidak’s): ***p<0.001. Comparisons are to wild-type donors. **(E,F)** Anatomical recovery of transplanted *tcima/b*^-/-^ mutant hosts was measured by anterograde axon labeling (E) or glial bridging (F). Sample sizes (axon regrowth): Vehicle (24), wild-type donor (21), *tcima/b*^-/-^ donor (19). Sample sizes (glial bridging): Vehicle (24), wild-type donor (20), *tcima/b*^-/-^ donor (23). One-way ANOVA (Sidak’s): ***p<0.001. Comparisons are to wild-type donors.

To examine whether *tcima/b* mutants elicit a cell-autonomous immune defect that impairs SC repair, we transplanted wild-type and *tcima/b^-/-^* kidney cells into *tcima/b^-/-^* fish, performed SC transections on transplanted animals at 14 dpt, and assessed swim endurance and axon regrowth at 28 dpi (which corresponds to 42 dpt). Compared to either vehicle injections or *tcima/b^-/-^* hematopoietic transplantation, wild-type kidney transplants enhanced swim endurance by 87.6% (p<0.0001) and proximal axon regrowth by 149.4% (p<0.0001) in *tcima/b^-/-^*hosts (Fig. 3D-E). By Gfap immunostaining, glial bridging was not statistically different across cohorts (Fig. 3F). Thus, underscoring an immune requirement for Tcim, transplanting wild-type leukocytes is sufficient to rescue the regeneration defects in *tcima/b* mutants.

### Tcim directs microglia/macrophage-mediated phagocytosis during SC regeneration

To determine putative binding partners of zebrafish Tcima, we next performed proteomics on zebrafish lysates. To this end, we immunoprecipitated recombinant, GST-tagged Tcima (Fig. S3A) with protein lysates collected from either whole zebrafish larvae at 2 days post-fertilization or adult SC tissues at 3 dpi (Fig. S3B). Trapped ion mobility (timsTOF) mass spectrometry of immunoprecipitated protein complexes identified 322 putative Tcima binding partners, including 59 proteins that were detected in both lysates, in addition to Tcima (Fig. 4A-B, S3C and Table S6). Our Tcima interactome enriched for genes implicated in cytoskeletal organization, fatty acid degradation, the complement pathway, and phago/endocytosis (phagosome) (Fig. 4C-D and Table S7). These results suggested Tcima may regulate lipid processing and/or phagocytosis during SC regeneration.

**Figure 4.**
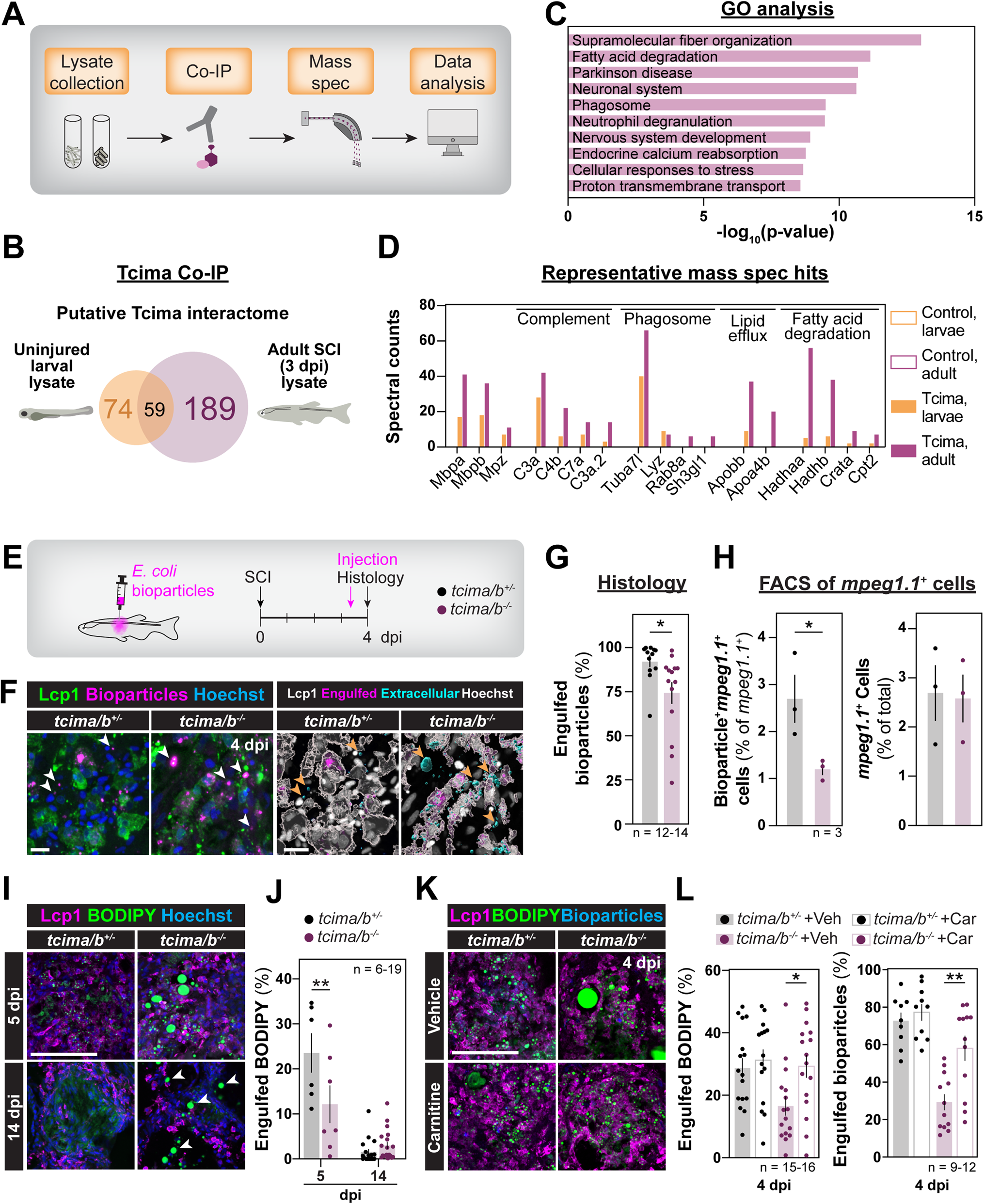
Tcim directs the phagocytic capacity of microglia/macrophages during SC regeneration. **(A)** Recombinant GST-Tcima was used for immunoprecipitation (Co-IP) with whole larvae or injured SC protein lysates. Co-IP was followed by timsTOF mass spectrometry. **(B,C)** Putative Tcima interacting proteins were identified from whole 2 dpf larvae or 3 dpi adult SC (B), and analyzed for GO term enrichment (C). **(D)** Spectral counts for top hits from (C) and other representative categories (including lipid efflux and the complement pathway). **(E)** *in vivo* phagocytosis assays in *tcima/b^-/-^* animals and control siblings. AlexaFluor-conjugated E. coli bioparticles were injected into the lesion site at 3 dpi. Fish were collected 16 hours after bioparticle injection for histology. **(F,G)** Histology of bioparticle-injected *tcima/b^-/-^* and control fish. Imaris was used to partition bioparticles into Extracellular (cyan) or Engulfed (magenta) channels based on Lcp1 surfaces (gray). Scale bars, 10 µm. Sample sizes: *tcima/b*^+/-^ (12) and *tcima/b*^-/-^ (14). Unpaired t-test (Welch’s): *p<0.05. **(H)** FACS of *Tg(mpeg1.1:YFP)*; *tcima/b*^-/-^ and *Tg(mpeg1.1:YFP)*; *tcima/b*^+/-^ fish that were injected with AlexaFluor-conjugated E. coli bioparticles at 3 dpi. The fraction of *mpeg1.1*:YFP^+^ cells that contained bioparticle fluorescence (left) and fraction of all *mpeg1.1*:YFP^+^ live cells (right) are shown. Samples represent 4-5 SCs per replicate and 3 replicates were used for each genotype. Unpaired t-test: *p<0.05. **(I,J)** BODIPY and Lcp1 staining in *tcima/b^-/-^* animals and control siblings at 5 and 14 dpi. Scale bar, 100 µm. Sample sizes: *tcima/b*^+/-^ 5 dpi (6), *tcima/b*^-/-^ 5 dpi (7), *tcima/b*^+/-^ 14 dpi (18), and *tcima/b*^-/-^ 14 dpi (19). One-way ANOVA (Sidak’s): **p<0.01. **(K,L)** Histology of carnitine-or vehicle-injected *tcima/b^-/-^* and control fish. All animals were injected with fluorescently-labeled bioparticles and stained with BODIPY and Lcp1. Scale bar, 100 µm. Sample sizes (BODIPY): *tcima/b*^+/-^ vehicle (16), *tcima/b*^+/-^ L-carnitine (15), *tcima/b*^-/-^ vehicle (16), and *tcima/b*^-/-^ L-carnitine (16). Sample sizes (bioparticle): *tcima/b*^+/-^ vehicle (9), *tcima/b*^+/-^ L-carnitine (10), *tcima/b*^-/-^ vehicle (12), and *tcima/b*^-/-^ L-carnitine (12). Two-way ANOVA (Tukey’s): *p<0.05, **p<0.01.

To determine if phagocytosis of cellular debris is altered in *tcima/b* mutants, we first measured lesion-associated myelin debris by Mbp staining. At 14 dpi, myelin engulfment, quantified as Mbp^+^ punctae in Lcp1^+^ cells, decreased from 56.0% ± 7.5% in controls to 29.5% ± 6.5% in *tcima/b* mutants (p=0.041) (Fig. S3D-E). To confirm these results, fluorescently-labeled heat-inactivated E. coli bioparticles were injected into the lesion at 3 dpi, and animals were collected 16 hours after injection to measure the engulfment capacity of Lcp1^+^ cells (Fig. 4E). Relative to control Lcp1^+^ leukocytes, which engulfed 92.1% ± 3.2% of injected bioparticles within 16 hours of intralesional injection, *tcima/b* mutant leukocytes engulfed significantly fewer bioparticles (74.3% ± 6.2%; p=0.019) (Fig. 4F-G). By FACS-sorting, while the numbers of *mpeg1.1*^+^ cells did not change across genotypes, the proportions of bioparticle-containing *mpeg1.1*^+^ cells were reduced by 55.6% (p=0.0461) in *tcima/b* mutants compared to controls (Fig. 4H and S3F). These studies showed reduced phagocytic capacity in *tcima/b* mutants.

### Tcim directs lipid processing and fatty acid oxidation during SC regeneration

To examine whether lipid processing is defective in *tcima/b* mutant leukocytes, we stained neutral lipids using BODIPY dye. In control animals, lipid droplets were readily seen in amoeboid Lcp1^+^ cells at 5 dpi, but were drastically reduced at 14 dpi, indicating efficient lipid processing and clearance of foamy leukocytes under control conditions (Fig. 4I and S3G-H). Conversely, the engulfment of BODIPY^+^ droplets was reduced by 48.4% (p=0.0025) in *tcima/b* mutants at 5 dpi (Fig. 4I-J). At 14 dpi, *tcima/b* mutants showed persistent lipid droplet-containing Lcp1^+^ cells, in addition to large BODIPY^+^ liposomes in the extracellular space (Fig. 4I and S3G-H). The persistence of lipid-laden leukocytes and extracellular lipids in *tcima/b* mutants suggested that intracellular lipid clearance is compromised in the absence of Tcim, and that stimulating lipid metabolism would rescue the phagocytic and lipid defects in *tcima/b* mutants.

To stimulate lipid catabolism via fatty acid oxidation, we injected L-carnitine into the lesion site *tcima/b* mutant and control siblings at 3 dpi and collected SC tissues for histology 16 hours after treatment. L-carnitine stimulates fatty acid oxidation by shuttling long chain fatty acids into mitochondria for catabolism (Loving and Bruce, 2020). L-carnitine supplementation increases functional recovery and tissue sparing in rodent SCI models (Ewan and Hagg, 2016; Karalija et al., 2012; Patel et al., 2010), although the molecular mechanisms driving these changes are not completely understood. At 4 dpi and in control animals, total BODIPY^+^ area decreased by 50.6% (p=0.0185) following L-carnitine injection compared to vehicle-injected controls (Fig. 4K-L and S3I-J). Whereas vehicle-treated *tcima/b* mutants accumulated large extracellular lipid deposits, L-carnitine treatment normalized Bioparticle engulfment and lipid distribution in *tcima/b* mutants (Fig. 4K-L and S3I-J). In an *in vivo* phagocytosis assay, bioparticle clearance was 2-fold improved in L-carnitine-injected *tcima/b* mutants relative to vehicle treatment (Fig. 4K-L and S3J). These results indicated that stimulating fatty acid oxidation is sufficient to rescue phagocytosis and lipid processing defects in *tcima/b* mutants.

### Human *TCIM* enhances lipid processing, phagocytosis and SC regeneration in a microglia/macrophage-dependent manner

The amino acid composition of zebrafish Tcima is 56% identical to human TCIM (Fig. 5A). To examine putative regenerative functions of TCIM, we expressed human TCIM under control of a heat-inducible promoter to generate *Tg(hsp70l:hTCIM-2A-EGFP)* fish, and assessed the outcomes of TCIM expression on functional and anatomical SC repair (Fig. 5B and S4A) (Halloran et al., 2000). *Tg(hsp70l:hTCIM-2A-EGFP)* zebrafish and wild-type siblings were subjected to SC transections or sham injuries followed by daily heat shocks starting at 2 dpi (Fig. 5B). Swim endurance was comparable between sham-injured TCIM-expressing and wild-type siblings after 6 weeks of daily heat shocks, indicating TCIM expression did not alter swim endurance capacity in the absence of SCI (Fig. 5C). However, at 14 dpi, swim endurance of TCIM-expressing fish was significantly increased compared to controls, and comparable to the performance of control fish at 42 dpi (Fig. 5C). TCIM-expressing fish displayed a 114.5% (p=0.0349) increase in proximal axon regrowth, while glial bridging was not significantly increased compared to controls at 14 dpi (Fig. 5D-E). These results indicated that expression of human TCIM is sufficient to accelerate axon regrowth and functional recovery after SCI.

**Figure 5.**
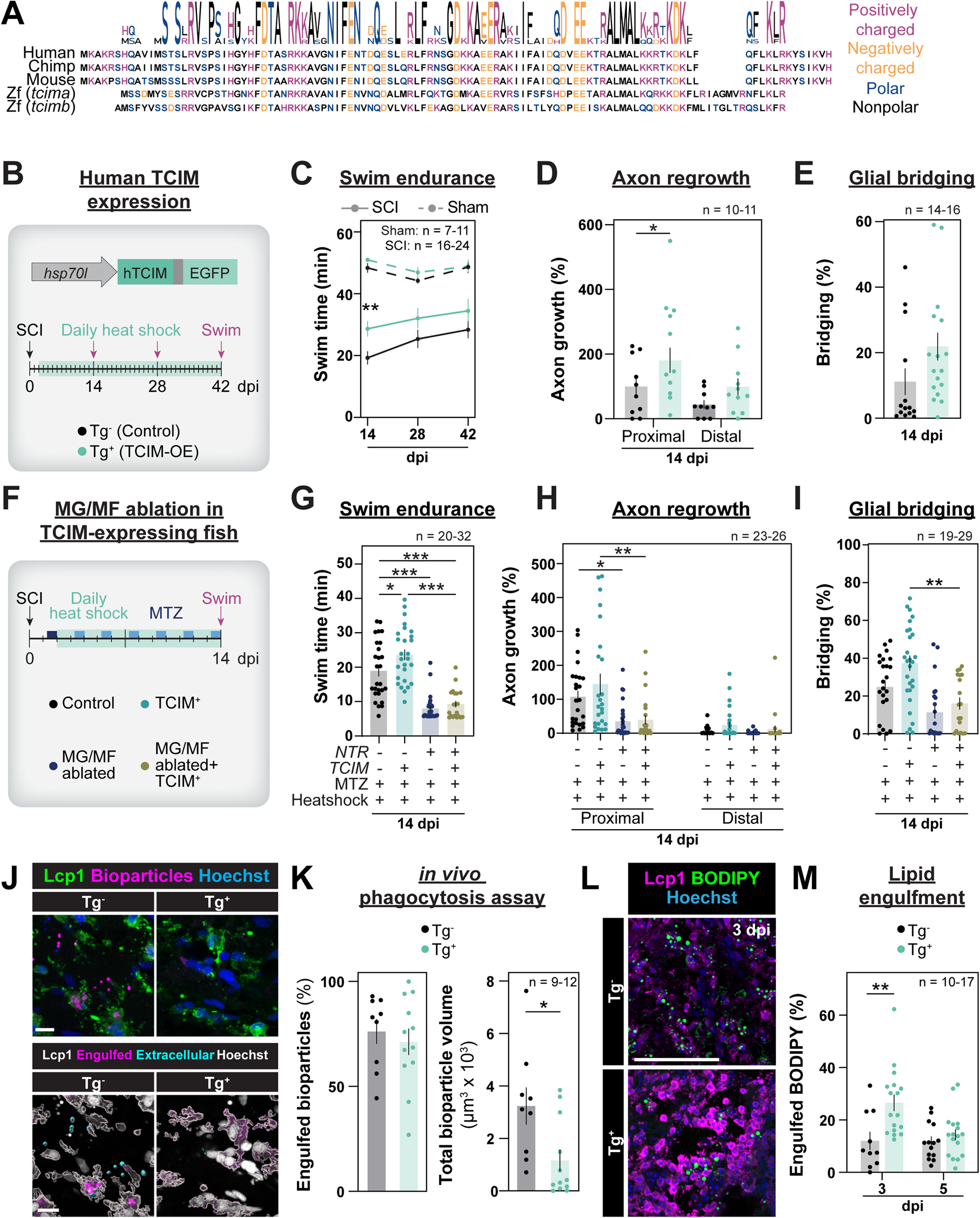
Expression of human TCIM enhances SC regeneration. **(A)** Sequence alignment and Biologo for human, chimpanzee, mouse, and zebrafish Tcim. **(B)** *Tg(hsp70l:hTCIM-2A-EGFP)* fish were generated for inducible expression of human TCIM. TCIM-expressing transgene-positive (Tg^+^) and control transgene-negative (Tg^-^) siblings were subjected to daily heat shocks starting at 2 dpi. Swim endurance was assessed at 14, 28, and 42 dpi and anatomical regeneration was assessed at 14 dpi. **(C)** Swim endurance of TCIM-expressing fish and control siblings at 14, 28, and 42 dpi. Dotted lines indicate sham-injured controls. Sample sizes: Control Sham (11), TCIM-expressing Sham (7), Control SCI (24), TCIM-expressing SCI (16). One-way ANOVA (Sidak’s): **p<0.01. Comparisons are to time-matched transgene-negative controls. **(D,E)** Anatomical regeneration in TCIM-expressing fish and control siblings was measured by anterograde axon labeling (D) or glial bridging (E). Sample sizes (axon regrowth): Control (10) and TCIM-expressing (11). One-way ANOVA (axon regrowth assay): *p<0.05. Sample sizes (glial bridging): Control (14) and TCIM expressing (16). Unpaired t-test (glial bridging). Comparisons are to transgene-negative controls. **(F)** Microglia/macrophage depletion via *Tg(mpeg1.1:NTR-IRES-EGFP)* was combined with expression of human TCIM via *Tg(hsp70l:hTCIM-2A-EGFP)*. Control and experimental fish were subjected to MTZ treatments starting at 1 dpi and daily heat shocks starting at 2 dpi. 5mM MTZ was used at 1 dpi followed by pulsed 1 mM MTZ between 3 and 14 days. **(G)** Swim endurance in *Tg(mpeg1.1:NTR-IRES-EGFP); Tg(hsp70l:hTCIM-2A-EGFP)* and control fish at 14 dpi. Sample sizes: NTR^-^TCIM^-^ (27), NTR^+^TCIM^-^ (32), NTR^-^TCIM^+^ (27), and NTR^+^TCIM^+^ (20). Two-way ANOVA (Tukey’s): *p<0.05, ***p<0.001. **(H,I)** Anatomical recovery in *Tg(mpeg1.1:NTR-IRES-EGFP); Tg(hsp70l:hTCIM-2A-EGFP)* and control fish. Axon regrowth (H) was measured by anterograde axon labeling and glial bridging (I) was measured by Gfap staining. Sample sizes (axon regrowth): NTR^-^TCIM^-^ (25), NTR^+^TCIM^-^ (26), NTR^-^TCIM^+^ (25), and NTR^+^TCIM^+^ (23). Sample sizes (glial bridging): NTR^-^TCIM^-^ (27), NTR^+^TCIM^-^ (27), NTR^-^TCIM^+^ (29), and NTR^+^TCIM^+^ (19). Two-way ANOVA (Tukey’s): *p<0.05, **p<0.01. **(J,K)** *in vivo* phagocytosis assays in TCIM-expressing and transgene-negative controls. All fish were subjected to daily heat shocks starting at 2 dpi, bioparticle injection in the lesion at 3 dpi, and tissue collection 16 hours after injection. Scale bars, 10 µm. The proportions of bioparticles within Lcp1^+^ cells (engulfed, left) and total bioparticle volume (right) were quantified. Sample sizes: Control (9) and TCIM-expressing (12). Unpaired t-test (Welch’s): *p<0.05. **(L,M)** BODIPY and Lcp1 staining in TCIM-expressing and control fish at 3 and 5 dpi. Scale bar, 100 µm. Sample sizes: Control 3 dpi (10), TCIM-expressing 3 dpi (17), Control 5 dpi (14), TCIM-expressing 5 dpi (17). One-way ANOVA (Sidak’s): **p<0.01. Comparisons are between and TCIM-expressing fish and their time-matched control groups.

To determine if accelerated regeneration in TCIM-expressing animals requires microglia/macrophages, we crossed *Tg(hsp70l:hTCIM-2A-EGFP)* animals into the *Tg(mpeg1.1:NTR-IRES-EGFP)* background. Compound transgenic fish were treated with daily heat shock from 2 to 14 dpi to induce TCIM expression and with pulsed MTZ doses from 1 to 14 dpi to ablate microglia/macrophages (Fig. 5F). Similar to microglia/macrophage depletion in wild-type fish, microglia/macrophage depletion impaired functional and anatomical regeneration metrics in TCIM-expressing fish (p=0.89 for swim endurance, p=0.99 for proximal axon regrowth, p=0.80 for glial bridging) (Fig. 5G-I). These results indicated that human TCIM enhances SC repair via a microglia/macrophage-dependent mechanism.

To test whether TCIM accelerates SC repair by promoting phagocytic clearance, we performed an *in vivo* phagocytosis assay in TCIM-expressing fish after SCI. *Tg(hsp70l:hTCIM-2A-EGFP)* animals or wild-type siblings were subjected to SC transections and exposed to daily heat shocks at 2, 3, and 4 dpi. Fluorescently-labeled bioparticles were injected into the lesion site at 3 dpi and animals were collected 16 hours after injection. Compared to control siblings, TCIM expression reduced total bioparticle volume by 64.1% (p=0.0249) at 4 dpi (Fig. 5J-K), and enhanced lipid engulfment by 54.6% (p=0.0025) at 3 dpi (Fig. 5L-M). Thus, human TCIM enhances SC regeneration by accelerating the engulfment and clearance of lipid-rich debris following SCI.

### TCIM expression reprograms myeloid precursors after SCI

To elucidate the molecular mechanisms by which TCIM enhances phagocytic clearance, we performed single nuclear RNA-sequencing (snRNA-seq) on TCIM-expressing SC tissues. *Tg(hsp70l:hTCIM-2A-EGFP)* and control fish were subjected to SC transections, daily heat shocks, and nuclear collection at 5 dpi (Fig. 6A). Sequencing was performed on 2 biological replicates of 35-39 pooled SCs per replicate. A total of 58,973 nuclei were obtained for downstream analysis using the Seurat package (Fig. S4B-D). Clustering of SC nuclei revealed 38 cell clusters with distinct molecular identities (Fig. 6A). Cell identities were determined using our previously developed hypergeometric scoring tool that optimized cell identification for non-mammalian vertebrates (Fig. S4E) (Saraswathy *et al*., 2023). At the pan-neural level, TCIM-expressing fish and wild-type siblings showed similar cell composition and proportions (Fig. S4F-G). *tcima* was exclusively expressed in leukocytes, while *tcimb* showed lower expression in leukocytes and non-immune cells including the endothelium (Fig. S4H). We noted that leukocyte proportions were not altered in the complete dataset upon TCIM expression (Fig. S4G). However, by subclustering leukocytes into 30 distinct clusters (Fig. 6A), cluster 2 was predominantly composed of wild-type cells, whereas clusters 3, 5, 6, and 29 were more prominent upon TCIM expression (Fig. 6B and S5A-B). These results indicated dynamic changes in immune cell identities in TCIM-expressing zebrafish.

**Figure 6.**
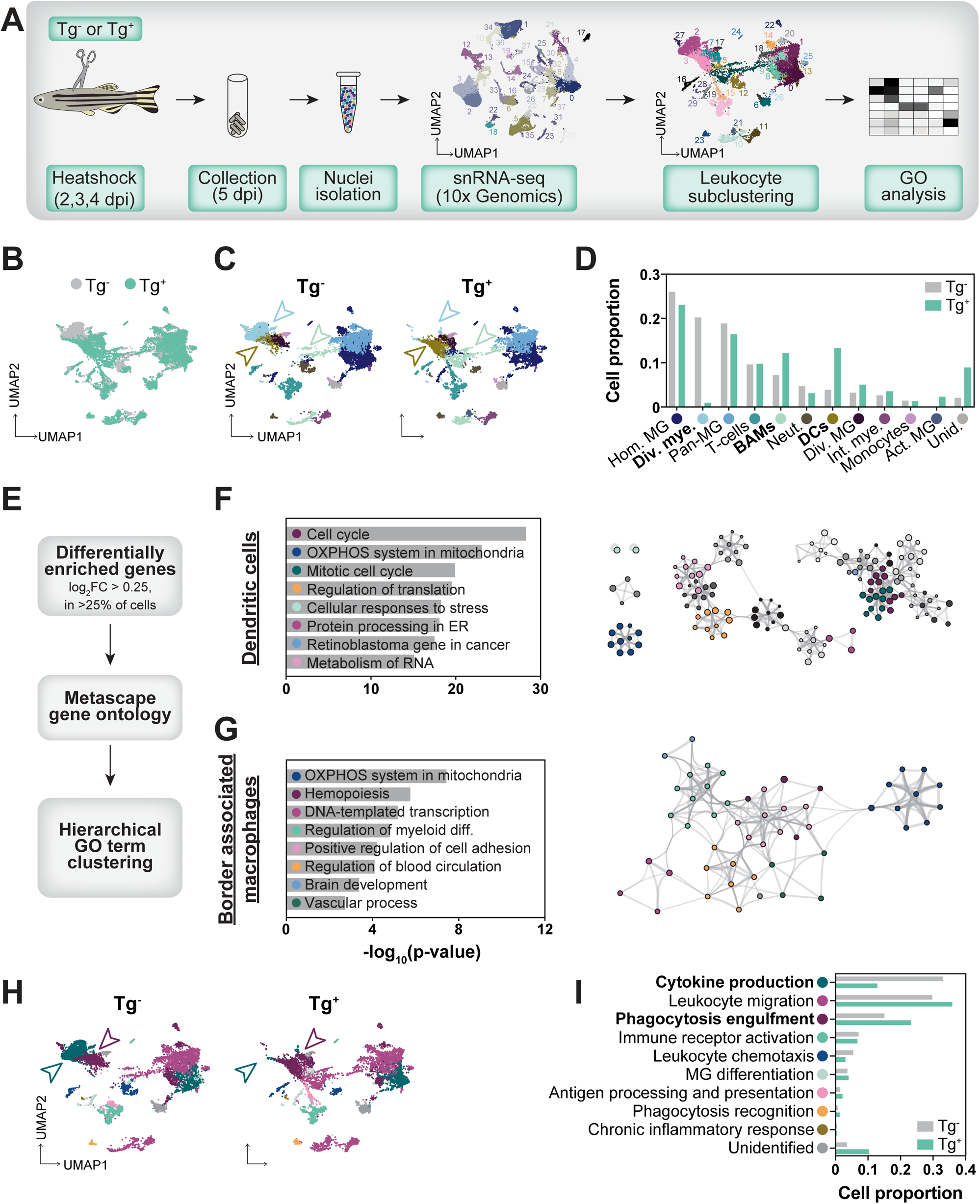
TCIM reprograms myeloid precursors into phagocytes to enhance SC repair. **(A)** snRNA-seq was performed on TCIM-expressing and control SC tissues at 5 dpi. All animals were subjected to daily heat shocks starting at 2 dpi. Sequencing was performed in biological duplicates and each duplicate represents 35-39 pooled SCs. Leukocytes were subclustered for further analysis. **(B,C)** UMAP representation of leukocyte subclusters in TCIM-expressing and control fish. Unbiased Seurat subclustering was used to identify immune cell identities. Subclusters that changed in proportion upon TCIM expression are indicated with arrowheads. **(D)** Proportions of the leukocyte subclusters shown in C. Leukocyte subtypes that changed upon TCIM expression are shown in bold. Homeostatic microglia (Hom. MG), dividing myeloid (Div. mye.), border associated macrophages (BAMs), neutrophils (Neut.), dendritic cells (DCs), dividing microglia (Div. MG), interferon myeloid (Int. mye.), activated microglia (Act. MG). **(E)** Top markers of leukocyte clusters that changed in proportion between TCIM-expressing and control siblings were analyzed for GO term enrichment. Top marker genes were expressed by >25% of cells within a cluster, enriched >0.25% log_2_(fold enrichment) from all other leukocyte subtypes, and had a p-value of <0.01 with the Wilcoxon Rank Sum test. **(F,G)** GO analysis was performed on top marker genes identified from the pipeline schematized in (E). Genes that were found as a top marker were used to produce GO term lists for dendritic cells (F), or border associated macrophages (G) subclusters. Metascape GO term clustering schematics are shown. Node size is proportional to the number of input genes falling into each GO term, and the line weight indicates the similarity in gene lists between connected GO terms. **(H)** UMAPs for leukocytes colored based on the most enriched immune-related GO term (from the immune GO term database located in Table S12) within the top marker genes from each subcluster. GO term subclusters that change in proportion upon TCIM expression are indicated with arrowheads. **(I)** Leukocyte proportions based on GO term subclustering. Leukocyte subtypes that change between TCIM-expressing and control fish are shown in bold.

To determine TCIM-induced cell identities, we performed hypergeometric scoring against two leukocyte marker databases (Fig. S5C) (Milich *et al*., 2021; Saraswathy *et al*., 2023). A complete list of genes used to identify leukocyte subtypes is shown in Table S8. This cell type classification identified 11 immune subtypes (homeostatic microglia, dividing myeloid, pan-microglia, T-cells, border associated macrophages, neutrophils, dendritic cells, dividing microglia, interferon myeloid, monocytes, and activated microglia) (Fig. 6C). We confirmed the expression of known zebrafish leukocyte markers, including pan-leukocytes (*lcp1*), myeloid cells (*ptprc*), microglia/macrophages (*mpeg1.1, csf1ra, and csf1rb*), microglia (*p2ry12, ccl34b.1, and lgals3bpb*), macrophages (*mfap4*), neutrophils (*mpx and lyz*), and T-cells (*lck and foxp3a*) (Fig. S5D). *tcima* was strongly enriched in microglia, while *tcimb* showed lower expression in a subset of leukocytes (Fig. S5D). The proportion of dividing myeloid cells was markedly reduced from 20.2% in controls to 0.97% in TCIM-expressing tissues, whereas two populations of mononuclear phagocytes, border associated macrophages (BAMs) (Green et al., 2022) and dendritic cells (Zhou et al., 2023), were 1.69 and 3.41-fold increased upon TCIM expression (Fig. 6C-D). These results suggested TCIM expression reprograms immature myeloid precursors into differentiated, phagocytic myeloid cells after SCI.

### TCIM expression alters myeloid gene expression toward lipid metabolism and debris clearance

To gain insights into the biological processes that are differentially regulated in TCIM-expressing SC tissues, we identified the top markers for the dividing myeloid (cluster 2), BAMs (clusters 5, 10, and 29), and dendritic (cluster 3) cell clusters for GO analysis (Fig. 6E). In support of their immature myeloid identity, cluster 2 cells enriched for genes related to cell division, transcription, and translation (ribosome, cell cycle, and transcriptional regulation by TP53) (Fig. S6A and Table S9). Consistent with their dendritic cell identity, cluster 3 cells were enriched for genes involved in antigen processing (which fell under the protein processing in ER GO term), including *ctsl* and *kif11* (Fig. 6F and Table S10). On the other hand, top markers of the BAMs clusters 5, 10, and 29 were implicated in vascular processes (regulation of cell adhesion, regulation of blood circulation, and vascular process in circulatory system), including expression of *itga4*, *tfrc*, *mef2a*, and *zfpm1* (Fig. 6G). Notably, genes related to phagocytic cup formation (*rab44* and *actg1*) were elevated, while negative regulators of lipid metabolism (*pde3b*, *zbtb20*, *rptor*, and *st3gal3*) were depleted in TCIM-expressing macrophages (Table S11). These findings supported a model in which phagocytes and lipid processing are enhanced upon TCIM expression.

To further explore the biological processes regulated by TCIM, we assembled a comprehensive list of genes related to “immune” or “leukocyte” GO terms using the Amigo gene ontology database (Table S12). By comparing this “immune GO term database” to the top markers of each leukocyte subcluster, we performed a hypergeometric probability test to identify the GO term that is most associated with each cluster (Fig. S6B). By pseudo coloring clusters based on their assigned GO term, the proportions of cytokine-producing leukocytes decreased by 61.3% upon TCIM expression (Fig. 6H-I). Conversely, the proportions of leukocytes that were enriched for “phagocytosis engulfment” genes increased by 55.3% in TCIM-expressing fish compared to controls (Fig. 6H-I). We also found “phagocytosis engulfment” gene expression was specifically enriched in TCIM-expressing leukocytes (Fig. S6C). These genes are largely involved in early steps of phagocytosis, such as formation of cell projections (*aif1l* and *cfl1*), phagocytic cup formation (*epcam*, *rab35b*, and *rac1a*), or phagocytic receptors (*scarb1*) (Fig. S6D-F). These results indicated human TCIM reprograms the molecular machinery of myeloid cells to enhance phagocytic capacity and SC regeneration.

## DISCUSSION

This study shows that phagocytic competence regulates differential regenerative capacity between mammals and zebrafish after spinal cord injury. By elucidating the cellular landscape and molecular identities of immune cells after SCI, we found that Tcim reprograms dividing myeloid cells into phagocytic macrophages to support efficient debris clearance and innate SC repair.

The fundamental principles that enable or disable regenerative capacity across species have eluded scientists for ages. By comparing the profiles of immune cells between highly regenerative zebrafish and less regenerative mammals, we identified species-specific immune features and mechanisms that direct innate SC repair. After mammalian SCI, pro-regenerative immune responses are overshadowed by prolonged and excessive activation of pro-inflammatory gene signatures that worsen injury outcomes. We show that, unlike mammals, adult zebrafish exhibit transient immune activation and efficient debris clearance. Consequently, zebrafish achieve innate SC repair in the absence of chronic inflammation. These findings are consistent with prior studies showing prolonged inflammation in macrophage-less zebrafish larvae (Tsarouchas *et al*., 2018). Although our profiling revealed dampened neutrophil and adaptive responses compared to macrophages, neutrophils are among the first immune cells to respond to SCI in zebrafish and mammals (Sena-Tomás et al., 2024; Taoka et al., 1997; Tsarouchas *et al*., 2018) and regulatory T-cell ablation impairs SC repair in zebrafish (Hui *et al*., 2017). Our comparative studies are consistent with zebrafish macrophages eliciting superior phagocytic capacity, which enables debris clearance and regeneration. Zebrafish possess myelin-associated proteins such as NogoA, but their inhibitory effects are thought to be dampened compared to mammals (Abdesselem et al., 2009). Our studies indicate that lipid-rich debris are cleared prior to axon regrowth and that manipulations that delay debris clearance impair regeneration. These findings suggest that enhanced phagocytosis, rather than differential inhibitory effects of myelin-associated debris, underscores elevated regenerative capacity of zebrafish. We propose that future comparisons and zebrafish studies will contribute new avenues to fine-tune immune SCI responses for neural repair.

The metabolic state of myeloid cells influences their cellular identity (Baseler et al., 2016; Jha et al., 2015; Kawamura et al., 2012; Vats et al., 2006). Our results support a model whereby zebrafish macrophages modulate their metabolic state and phagocytic output to support SC repair. Biomolecule catabolism is necessary for monocytes and myeloid precursors to differentiate into macrophages (Jacquel et al., 2012; Zhang et al., 2012), while adopting a phagocytic phenotype is sufficient to shift macrophages toward a reactive phenotype during muscle regeneration (Zhang et al., 2019). Lipid processing is a chief component of biomolecule catabolism. Lipid-laden myeloid cells that fail to process and recycle lipids transition into a hyper-inflammatory, foamy phenotype (Wang *et al*., 2015b). On the other hand, lipid breakdown and fatty acid oxidation enhance phagocytic capacity and promote a neuroprotective phenotype in microglia in mouse models of neurodegeneration (Leng et al., 2022). While limited lipid processing and foamy macrophages persist months to years after mammalian SCI (Fleming *et al*., 2006; Pruss *et al*., 2011), we found that lipid-rich *mpeg1.1*^+^ cells are transient and disappear by 14 dpi in zebrafish. We propose that efficient lipid processing stimulates a feed-forward loop that enhances phagocytic capacity in zebrafish macrophages.

Our data support a model whereby Tcim directs lipid processing and debris clearance to promote SC repair. Consistent with disrupted lipid homeostasis, lipid droplets and foam cells persist for weeks in *tcima/b* mutant SC tissues. Conversely, TCIM expression is sufficient to expand phagocytic macrophages at the expense of dividing myeloid cells. As lipid and biomolecule breakdown was shown to shift myeloid cells towards differentiation or activation (Zhang *et al*., 2019; Zhang *et al*., 2012), we propose that TCIM alters immune cell composition by enhancing leukocytic lipid catabolism. Our findings suggest TCIM function is conserved across vertebrates but that its expression is not. Murine Tcim is expressed by microglia and macrophages in the uninjured SC and at 1 dpi, but is downregulated at 7 dpi (Milich *et al*., 2021; Salvador et al., 2023), while human TCIM is not detectable in homeostatic microglia or macrophages (Zhang et al., 2016). Our Tcima interactome included multiple proteins in the lipid trafficking, catabolism, and recycling pathway. Consistent with Tcima acting directly in lipid metabolism, protein structure prediction software identified a putative intrinsically disordered region featuring a Membrane-binding Molecular Recognition Feature (MemMoRF) (Basu et al., 2023), which suggests interactions directly with lipids and/or lipid membranes. In depth studies are needed to pinpoint the mechanism by which Tcim and its intrinsically disordered region impacts lipid recognition, engulfment, or processing in zebrafish and mammals.

Our findings shed light on phagocytic debris clearance as a key target to promote neural regeneration. Debris accumulation and phagocytosis impact functional outcomes in models of neural injury (Akhmetzyanova et al., 2018; Ritzel et al., 2015), multiple sclerosis (Berghoff et al., 2021; Boven et al., 2006), stroke (Neher et al., 2013), and neurodegeneration (Keren-Shaul et al., 2017; Krasemann *et al*., 2017; Pluvinage et al., 2019). We propose that zebrafish present a valuable model and screening platform to devise pro-regenerative immune cell interventions for neural repair.

## ACKNOWLEDGMENTS

We thank S. Ackerman, V. Cavalli, J. Kipnis and L. Solnica-Krezel (Washington University) for discussion; S. Nandagopal, T. Tsai (Washington University) and S. Megason (Harvard) for the HCR probe design script; B. Appel (University of Colorado), K. Monk (Vollum Institute), M. Redd (University College London), M. Roh-Johnson (University of Utah), K. Tanner (NIH/NCI), and T. Tsai (Washington University) for reagents and zebrafish lines. We thank the Zebrafish Shared Resource for animal care, the Siteman Cancer Center Flow Cytometry Core, the Center for Cellular Imaging, the Genome Access Technology Center, the Small Animal Radiation Research Platform, and the Proteomics Shared Resource at Washington University. This research was supported by grants from the Washington University Center of Regenerative Medicine (to D.K.S.), the W.M. Keck Foundation (to D.K.S.), NIH NIBIB (T32 EB028092 to D.K.S), NIH NICHD (F32 HD107935 to D.K.S.), NIH NINDS (K99 NS133484 to D.K.S. and R01 NS123708 to M.H.M.), and the Washington University Hope Center for Neurological Disorders (to M.H.M).

## ETHICS DECLARATIONS

The authors declare no competing interests.

## MATERIALS AND METHODS

### Zebrafish care and husbandry

Adult zebrafish of the Tubingen strain were maintained at the Washington University Zebrafish Core Facility. All experiments were performed in compliance with institutional animal protocols. Male and female animals ∼4 months of age were used. Experimental fish and control siblings of similar size and equal sex distribution were used for all experiments. Spinal cord transection surgeries and regeneration analyses were completed in a blinded manner, and two to three independent experiments were performed using different clutches of animals. Previously published zebrafish lines used in the study are as follows: *Tg(mpeg1.1:YFP)*^w200^ (Roca and Ramakrishnan, 2013), *Tg(lyz:EGFP)*^nz117^(Hall *et al*., 2007), *Tg(lck:lck-EGFP)*^cz2^ (Langenau *et al*., 2004), *Tg(mpeg1.1:NTR-IRES-EGFP-CAAX)*^co57^ (Hughes and Appel, 2020), and *Tg(act2b:membrane-Citrine)*^hm30^ (Xiong et al., 2014). Newly constructed strains are described below.

*Generation of tcima and tcimb mutant zebrafish.* sgRNA was prepared by *in vitro* transcription of double-stranded deoxyoligonucleotide templates as described in (Gagnon et al., 2014). gRNA target sequences are as follows: *tcima* (GGCATGAAATCTAAACTAAC and GGACAAGGCATAATGTTTAC) and *tcimb* (GGAACGGTTAGTGAACGCCC and GGACAAATTCATGCTGATTA). sgRNA was transcribed from the annealed double-stranded oligonucleotide template using MEGAshortscript™ T7 transcription Kit (Invitrogen) according to the manufacturer’s instructions. Transcribed sgRNA was purified using RNA clean & concentrator-25 (ZYMO Research) and recovered in distilled water. For microinjection, 25-30 ng/µL sgRNA was gently mixed with 0.5 ug/µL Cas9 (PNA BIO cat# CP01), and 0.5 µl 0.25% phenol red solution and diluted with RNase-free water to a final volume of 4 µL. Prior to microinjection, the RNP complex solution was incubated at 37°C for 5 min and then placed at room temperature. ∼1 nL of 5μM sgRNA:Cas9 RNP complex was injected into the cytoplasm of one-cell stage embryos.

The following primers were used for genotyping: *tcima*_common_F: ggccaccagaatttctaaaaga, *tcima*_wt_R: ctcctcggctttcatgtctc, and *tcima*_mutant_R: tgacaaagtcagtcaatttaagaca, *tcimb*_common_F: ggcgacatcagtccgtattt, *tcimb*_wt_R: aacgccatgtctttctacgtg, *tcimb*_mutant_R: aagttcatgcttgcgtcaga. The full name of the *tcim* mutant lines are *tcima^stl661^*and *tcimb^stl662^*. *Generation of Tg(hsp70l:hTCIM-2A-EGFP) zebrafish.* The following forward primers were used to amplify human TCIM (hTCIM) cDNA: ClaI forward primer 5’-gatcATCGATcgccaccatgaaagcaaagcgaagcc-3’ and XhoI reverse primer 5’-gatcCTCGAGgtgaactttgatggaat-3’ where the uppercase sequences indicate restriction enzyme sites. The genomic fragments were subcloned into ClaI/XhoI-digested pCS2-hsp70l:2A-EGFP plasmid. Clones were co-injected into one-cell stage wild-type embryos with I-SceI. Five founders were isolated and propagated for this transgene. The highest expressing line was selected for our studies. The full name of this line is *Tg(hsp70l:hTCIM-2A-EGFP)^stl682^*. Animals were analyzed as hemizygotes.

*Spinal cord transections and treatments.* Zebrafish were anaesthetized in 0.2 g/L of MS-222 buffered to pH 7.0. Fine scissors were used to make a small incision that transects the spinal cord 4 mm caudal to the brainstem region. Complete transection was visually confirmed at the time of surgery. Injured animals were also assessed at 2 and 3 dpi to confirm loss of swim capacity post-surgery.

*Heat shock treatments.* Zebrafish were maintained in a circulating system retrofitted with a controlled heater (Tecniplast). Water temperature was increased to 37 °C for 1 hour per day and gradually returned to 28 °C to induce transgene expression.

### Histology

Sixteen μm cross or 20 μm longitudinal cryosections of paraformaldehyde-fixed SC tissues were used. Serial sections were collected -150 µm apart for ∼3 mm rostral and caudal to the transection site.

For immunohistochemistry, tissue sections were circumscribed with a hydrophobic barrier pen and rehydrated in PBT (0.1% Tween-20 in PBS). After 2 x 5 min washes in PBT, sections were treated with blocking agent (5% goat serum in PBT) for 1 hour at room temperature. Sections were incubated overnight with indicated primary antibodies diluted in blocking agent, washed in PBT, and treated for 1 hour in secondary antibodies diluted in blocking agent. Following washes, sections were incubated in 1 μg/mL of Hoechst, washed in PBS, and mounted in Fluoromount-G (SouthernBiotech cat# 0100-01) mounting media. Primary antibodies used in this study were chicken anti-GFP (AVES, cat# 1020, 1:1000), mouse anti-Gfap (ZIRC, cat# Zrf1, 1:1000), rabbit anti-Lcp1 (a kind gift from Michael Redd, 1:10000), and rabbit anti-Mbp (a kind gift from Bruce Appel, 1:1000). Secondary antibodies (Invitrogen, 1:250) used in this study were AlexaFluor-488, AlexaFluor-594, and AlexaFluor-647 goat anti-rabbit, anti-mouse, or anti-chicken antibodies.

BODIPY 493/503 (Fisher Scientific cat# NC1707119) was reconstituted as a 1 mg/mL stock, which was diluted 1:250 in PBT at the time of use, and sections were incubated for 30 minutes in diluted BODIPY prior to several washes with PBT.

Hybridization Chain Reaction (HCR) RNA *in situ* hybridization protocol was adapted from Molecular Instruments (https://www.molecularinstruments.com/hcr-ihc-protocols) (Choi et al., 2018). HCR RNA probes used in this study (*tcima* and *tcimb*) were designed using a custom python script and ordered as 50 pmol opools (IDT). Oligo sequences are provided in Tables S13 and S14. Briefly, tissue sections were hydrated in PBS, dehydrated into 100% ethanol, then rehydrated stepwise into PBT (0.1% Tween-20 in PBS). Sections were boiled in citrate buffer (10mM Citric Acid, 0.05% Tween-20, pH 6.0) for 10 minutes, allowed to cool to room temperature, and then washed thoroughly in PBT. For blocking, sections were incubated in prewarmed hybridization buffer (Molecular Instruments) for 1 hr at 37°C. For hybridization, sections were incubated in prewarmed DNA probe sets diluted to 0.015 pmol/µL in hybridization buffer for 48 hrs at 37°C. Washes were performed in wash buffer (Molecular Instruments) at 37°C followed by 5x SSCT (3M NaCl, 0.3 M Sodium Citrate, 0.1% Tween-20, pH 7.0) at room temperature. For pre-amplification, sections were incubated in amplification buffer (Molecular Instruments) for 1 hr at room temperature. Prior to amplification, h1 and h2 hairpins were snap-cooled in individual tubes by heating to 95°C and allowing tubes to return to ambient temperature slowly. For amplification, snap-cooled hairpins were mixed and diluted 1:50 in Amplification buffer overnight at room temperature in the dark. Samples were washed in 5x SSCT, 5x SSC, and PBT at room temperature before proceeding with immunohistochemistry.

Tissue sections were imaged using a Zeiss AxioScan widefield microscope for axon regrowth and glial bridging, or a Zeiss LSM 800 confocal microscope for all other immunofluorescence. To image the lesion site, the imaging window was set immediately dorsal to the vertebral column. For immune histology, parenchymal sections 150 µm rostral (unless otherwise indicated) to the lesion were imaged.

### Fluorescence-Activated Cell Sorting (FACS)

Four mm SC tissue sections, including the lesion site plus additional rostral and caudal tissue proximal to the lesion, were collected from adult injured zebrafish or control uninjured siblings. Three biological replicates were used, and each replicate represents a pool of 4-5 SCs. Tissues were dissociated using 0.05% Trypsin for 5 min at room temperature and subjected to 3 washes with Hanks’ Balanced Salt Solution (HBSS, Gibco). Cell supernatants were triturated in DMEM media with 20% fetal bovine serum using a 1000 μL pipette and applied to a 100 μm cell strainer (MidSci, cat# 100ICS). Dissociated cells were pelleted at 500 g for 5 min, resuspended in HBSS with 2% FBS, and sorted using a MoFlo Cell sorter machine. FACS experiments were performed at the Siteman Cancer Center flow cytometry core. Wild-type or *Tg(mpeg1.1:YFP)* animals were used to set up YFP FACS gates. For the AlexaFluor-conjugated bioparticle phagocytosis assays, wild-type animals were injected and used to set gates for AlexaFluor-594. LIVE/DEAD cell stain (ThermoFisher cat# L34973) was used to label live cells. Sorted cells were collected into serum-supplemented DMEM for RNA extraction.

### Drug treatments

For metronidazole (MTZ) treatments, MTZ (Sigma cat# M1547-25G) was dissolved directly into fish water to a concentration of either 5 mM (1 dpi treatments), 1 mM (pulsed treatments), or 10 mM (transplant experiments). Fish were placed in individual treatment beakers. 5 mM treatments lasted 18 hours, while 1 mM or 10 mM treatments were for 24 hours, in the dark. MTZ treatments were pulsed every other day intermingled with fresh fish water treatments. In the case of combining MTZ treatment with daily heat shocks, MTZ treatments were performed for 18 hours in between daily heat shocks at standard water temperature (28 °C). The dosing and timing of drug treatments were optimized prior to performing experiments, and effective depletion was confirmed using histology for Lcp1. In our hands, we did not observe EGFP expression from the IRES-EGFP cassette present in *Tg(mpeg1.1:NTR-IRES-EGFP)* fish.

For L-carnitine treatments, 100 mg/mL (2x) stock of L-carnitine (Sigma cat# 8400920025) was made in sterile water. On the day of injection, this stock was diluted to 50 mg/mL with 1 mg/mL Escherichia coli bioparticles AlexaFluor-594 conjugate (ThermoFisher cat# E23370) as described in the phagocytosis assay section. Fish were placed upright in a slotted cellulose sponge and 5 µL of working solution was injected directly into the lesion site at 3 dpi with a Hamilton gastight syringe (cat# 80008). Dosing was determined based on previous administration of L-carnitine in zebrafish (Li et al., 2017).

### Swim endurance assays

Zebrafish were exercised in groups of 8-15 in a 5L swim tunnel device (Loligo, cat# SW100605L, 120V/60Hz). After 15 minutes of acclimation (5 minutes with no flow, 5 minutes with 9 cm/second flow, then 5 minutes with 10 cm/second flow) inside the enclosed tunnel, water current velocity was increased every two minutes and fish swam against the current until they reached exhaustion as described in (Burris et al., 2021). Exhausted animals were removed from the chamber without disturbing the remaining fish. Swim times at exhaustion were recorded.

### Anatomical regeneration assays

Anterograde axon tracing was performed on adult fish. Fish were anaesthetized using MS-222 and fine scissors were used to transect the cord 4 mm rostral to the lesion site. Biocytin-soaked Gelfoam Gelatin Sponge was applied at the new injury site (Gelfoam, Pfizer, cat# 09-0315-08; Biocytin, saturated solution, Sigma, cat# B4261). Fish were euthanized 4 hours post-treatment and biocytin was histologically detected using AlexaFluor-594-conjugated Streptavidin (Molecular Probes, cat# S-11227).

To calculate axon regrowth, biocytin labeled axons were quantified using the threshold and particle analysis tool in Fiji. Multiple sections for each fish at each level were quantified: +0.5 mm (caudal-proximal, four sections), +2 mm (caudal-distal, four sections), -1 mm (rostral, two sections). The number of biocytin^+^ particles in each region of the spinal cord were averaged for each individual fish. The caudal-proximal and caudal-distal values were divided by the rostral value to give the axon growth index. This value was then normalized to the average caudal-proximal value in sibling controls. To calculate glial bridging, the cross-sectional Gfap^+^ area of the center of the lesion was normalized the cross-sectional area of the spinal cord 1 mm rostral to the lesion site.

### Bulk RNA-sequencing

Total RNA was prepared using NucleoSpin® RNA Plus XS (Clontech, cat# 740990). RNA was prepared from 4 mm SC sections at 5 dpi in *tcima/b*^-/-^ and wild-type controls for bulk RNA sequencing. 9-10 SCs were pooled for each replicate and 3 biological replicates were used. RNA sequencing was performed at the Genome Technology Access Center at Washington University. Total RNA integrity was determined using Agilent Bioanalyzer or 4200 Tapestation. Library preparation was performed with 10 ng of total RNA with a Bioanalyzer RIN score greater than 8.0. ds-cDNA was prepared using the SMARTer Ultra Low RNA kit for Illumina Sequencing (Takara-Clontech cat# 634936) per manufacturer’s protocol. cDNA was fragmented using a Covaris E220 sonicator using peak incident power 18, duty factor 20%, cycles per burst 50 for 120 seconds. cDNA was blunt ended, had an A base added to the 3’ ends, and had Illumina sequencing adapters ligated to the ends. Ligated fragments were then amplified for 12-15 cycles using primers incorporating unique dual index tags. Fragments were sequenced on an Illumina NovaSeq-6000 using paired end reads extending 150 bases. All Metadata, raw data, and processed data associated with bulk RNA-sequencing can be found in the NCBI Gene Expression Omnibus.

### Gene ontology analysis

Gene ontology (GO) analysis was performed using Metascape (https://metascape.org/). Default parameters were used to cluster overlapping GO terms (minimum overlap of 3 genes, p-value cutoff of 0.01, minimum gene enrichment of 1.5). Gene sets selected for analysis were GO Biological Processes and KEGG Pathways. GO term titles were shortened in some cases for brevity in figures. The full list of term titles and hierarchical GO terms can be found in supplemental tables.

### Identification of pro-regenerative immune genes by single nuclear RNA-sequencing

A snRNA-seq atlas of the regenerating zebrafish SC was used (Saraswathy *et al*., 2023). To identify top markers of microglia/macrophages, we filtered for genes that were expressed in >10% of the cells identified in the microglia/macrophage 1 cluster, enriched with a log_2_(fold enrichment) >0.25 compared to all non-microglia/macrophage cells, and possessed a p-value of <0.01 with the Wilcoxon Rank Sum test. Genes upregulated (>0.25 log_2_(fold change)) at 7 dpi compared to uninjured were selected. To identify injury-induced mouse microglia marker genes, a single cell RNA-sequencing atlas of mouse SCI was used (Milich *et al*., 2021). Selected top markers showed >10% cell expression in microglia, log_2_(fold enrichment) >0.25 compared to all other cell clusters, and log_2_(fold change) >0.25 after SCI (either up or down).

### Isolation of spinal cord nuclei for RNA sequencing

Nuclei were isolated as previously described (Saraswathy *et al*., 2023). Three mm SC tissue, flanking the lesion sites were collected from 35-39 adult zebrafish at 5 dpi from heat shocked (at 2, 3, and 4 dpi) *Tg(hsp70:hTCIM-2A-EGFP)* and wild-type siblings. For tissue lysis, SC tissues were homogenized at a low setting for 15 seconds. Density gradient separation using sucrose solution was used to sediment nuclei from the supernatant. Final nuclear lysates were resuspended using 100 µl of resuspension solution (1x PBS, 2% BSA, 0.2U/µl RNase inhibitor; New England Biolabs, cat# M0314S). Hoechst staining was performed to assess the quality of isolated nuclei based on their morphology. Samples in which more than 70% of the nuclei were scored as ‘healthy’ were submitted for snRNA-seq.

### Single nuclear RNA sequencing

For snRNA-seq, 30 µl of isolated nuclei at a concentration of ∼1000 nuclei/µl was submitted to Genome Technology Access Center at McDonnel Genome Institute of Washington University. Two biological replicates of each genotype were used. cDNA was prepared after GEM generation and barcoding, followed by the GEM-RT reaction and bead clean-up steps. cDNA was amplified for 11-13 cycles and purified using SPRIselect beads. Purified cDNA samples were run on a Bioanalyzer to determine the cDNA concentration. GEX libraries were prepared as recommended by the 10x Genomics Chromium Single Cell 3’ Reagent Kits User Guide (v3.1 Chemistry Dual Index) with appropriate modifications to the PCR cycles based on the calculated cDNA concentration. For sample preparation on the 10x Genomics platform, the Chromium Next GEM Single Cell 3’ Kit v3.1, 16 rxns (PN-1000268), Chromium Next GEM Chip G Single Cell Kit, 48 rxns (PN-1000120), and Dual Index Kit TT Set A, 96 rxns (PN-1000215) were used. The concentration of each library was accurately determined through qPCR utilizing the KAPA library Quantification Kit according to the manufacturer’s protocol (KAPA Biosystems/Roche cat# KK4824) to produce cluster counts appropriate for the Illumina NovaSeq6000 instrument. Normalized libraries were sequenced on a NovaSeq6000 S4 Flow Cell using the XP workflow and a 50x10x16x150 sequencing recipe according to manufacturer protocol. A median sequencing depth of 50,000 reads/cell was targeted for each Gene Expression Library. All Metadata, raw data, and processed data associated with snRNA-sequencing from this study can be found in the NCBI Gene Expression Omnibus.

### Aligning snRNA-seq reads

After sequencing, the Illumina output was processed using the CellRanger (v6.0.0) recommended pipeline to generate gene-barcode count matrices. A custom reference genome was generated with the “cellranger mkref” command using the fasta file of zebrafish reference genome GRCz11 constructed from the Ensemble genome build (https://useast.ensembl.org/Danio_rerio/Info/Index) and the sorted Gene Transfer Format file (v4.3.2) from the improved zebrafish transcriptome annotation (Lawson et al., 2020). Base call files for each sample from Illumina were demultiplexed into FASTQ reads. Then, the “cellranger count” pipeline was used to align sequencing reads in FASTQ files to the custom reference genome. Both exon and intron sequences were aligned. The filtered gene-barcode count matrices generated by “cellranger count” was used for downstream analysis.

### Integrated analysis of snRNA-seq data

All the datasets were combined and analyzed using Seurat (v4.1.1) package with R (v4.2.1) (R Core Team, 2018; (Stuart et al., 2019)). Each sample count matrix was filtered for genes that were expressed in at least 3 cells and cells expressing at least 200 genes, followed by cell quality assessment using commonly used QC matrixes (Ilicic et al., 2016). Cells having a unique number of genes between 200 to 4000 and a mitochondrial gene percentage <5 were used for downstream processing. Each dataset was independently normalized and scaled using the “SCTransform” function, which is an improved method for normalization, that performs a variance-stabilizing transformation using negative binomial regression (Hafemeister and Satija, 2019). Standard integration workflow of Seurat was used to identify shared sources of variation across experiments as well as mutual nearest neighbors (Butler et al., 2018; Haghverdi et al., 2018). Integration features were selected based on the top 4000 highly variable features using “SelectIntegrationFeatures” function (nfeatures = 4000), which was used as input for the “anchor.features” argument of the “FindIntegrationAnchors” function. Principle Component Analysis (PCA) analysis was performed on the 4000 variable features and the top 50 principal components selected based on the elbow plot heuristic, which measures the contribution of variation in each component. These 50 principal components were used in “FindNeighbors” and “FindClusters” functions to perform graph-based clustering on a shared nearest neighbor graph (Levine et al., 2015; Xu and Su, 2015). Louvain algorithm was used for modularity optimization in clustering the cells using “FindClusters” function. The resolution parameter (res = 0.5) that determines the granularity of the clustering was selected by visually inspecting clusters with resolutions ranging 0.1 - 2.0 as well as clustree graphs (Zappia and Oshlack, 2018). Uniform Manifold Approximation and Reduction (UMAP) was used for non-linear dimensional reduction of the first 50 principal components and visualize the data using “RunUMAP” function (Becht et al., 2018). Data was graphed using different plot functions, such as “DimPlot”, “VlnPlot”, “FeaturePlot”, “Dotplot” and “DoHeatmap”, to view the cell cluster identity and marker gene expression. Cell proportion data was extracted using “table” and “prop.table” functions. Differential gene expression for individual cluster was identified using Wilcoxon rank sum test in the “FindAllMarkers” function. Marker genes detected in at least 25% of the clustered cells and with a log_2_(fold change) threshold of 0.25 were selected, unless mentioned otherwise. Only positive markers were reported.

### Subset analysis of leukocyte clusters

Leukocytes identified from the complete dataset were subclustered using the “subset” function for subcluster analysis. The subset was normalized and scaled using “SCTransform” function with glmGamPoi method (Ahlmann-Eltze and Huber, 2021). Fifty principal components were used, and the resolution parameter was set to 0.7. Further downstream analysis was done as described above for the integrated analysis.

### Cluster identification using differentially expressed markers

Cell identities were determined as previously described (Saraswathy *et al*., 2023). Briefly, a “<u>V</u>ertebrate C<u>N</u>S <u>M</u>arkers” database of previously published cell markers of vertebrate brain and/or SC tissues was compiled (Baek et al., 2019; Cavone *et al*., 2021; Guillemot, 2007; Haring et al., 2018; Hayashi et al., 2018; Hernandez-Miranda et al., 2017; Lu et al., 2015; Milich *et al*., 2021; Rosenberg et al., 2018; Rougeot et al., 2019; Sathyamurthy et al., 2018; Tambalo et al., 2020; Tang et al., 2017; Yadav et al., 2023; Zeisel et al., 2018; Zhang et al., 2014). For each cell cluster, every marker gene identified a top differentially expressed (DE) marker of that cluster was cross-referenced with our compiled VNM database using our “DEMarkerScoring” algorithm. For every matching marker gene, one point was given to the respective cluster under the column name with the matching cell identity. Iteration over every marker gene was performed to generate a scoring matrix with varying points for each cluster against the different cell identities compiled in the VNM database. The “phyper” function in R was then used to calculate binomial probabilities using hypergeometric distribution for the total score obtained by each cluster against each cell identity in the database. The -log_10_ of probability values were then calculated. The resulting values were scaled to 0-100 and plotted as a heatmap using GraphPad Prism. Each cluster was given an identity based on the maximum -log_10_ P score obtained in the heatmap. The top DE markers of clusters with ambiguous scores were manually searched in the literature (Zhang *et al*., 2014; Zhang *et al*., 2016) to confirm cluster identity. “RenameIdents” function was used to assign identity to each cluster. For each cluster, cell identities were confirmed by surveying canonical marker expression. A similar method was used for subclustering leukocytes, but an altered “Leukocyte markers” database that contained leukocyte identities in the VNM database was used. A full list of “Leukocyte markers” are detailed in Table S8.

For gene ontology-based cell clustering, the GO term database Amigo (http://amigo.geneontology.org/amigo/landing) was used. To identify immune-related biological processes, the Amigo Drilldown Viewer was first used to view GO term hierarchy. All terms located directly in the layer under “immune system process” were selected. Terms with 15 to 400 genes that aligned to the *Danio rerio* genome were used. Second, the keyword search terms “immune” and “leukocyte” were used to identify more specific immune-related GO terms. These GO terms were also filtered by selecting terms with 50-400 genes that aligned to the Danio rerio genome. Following filtering, prospective gene lists were analyzed by Metascape. Any gene list that did not return the GO term in question within the top three results was removed from the analysis. Only GO terms that were the most statistically significant amongst the leukocyte subclusters are shown. The full list of genes with this “Leukocyte GO term” database is in Table S12.

### Transplantation of whole kidney marrow

For transplantation experiments, *Tg(mpeg1.1:NTR-IRES-EGFP)* fish in the *tcima/b*^-/-^ background were treated for 24 hours with 10 mM MTZ as described in the drug treatment section. Adult zebrafish were irradiated with 8 Gy radiation in the xStrahl Small Animal Radiation Research Platform (SARRP) in the Radiology Department at Washington University for two consecutive days. Individual fish were placed in 5 cm petri dishes in fresh fish water on the radiator platform in a single layer for irradiation. After each radiation dose, individual fish were recovered in beakers in 100-150 mL of fresh system water. The day after the second radiation dose, fish received dissociated marrow cells or vehicle.

To collect cells for transplantation, kidney tissues were dissected from *Tg(actb2:membrane-Citrine)^hm30^* donor fish (in either a wild-type or *tcima/b*^-/-^ background) and placed in chilled HBSS+H (HBSS, 1% HEPES). Kidneys were treated with 0.05% Trypsin for 5 min at room temperature, and then washed with HBSS+H. Cells were dissociated with 30 brisk triturations in DMEM+20% FBS, placed through a 100 µm cell strainer (MIDSCI cat# ASCELLSTAIN2), and resuspended in HBSS+2% FBS. Phenol red was added (1:1000) in cell suspensions or in vehicle for injections. For each experiment, 250,000-350,000 donor cells or the same volume of empty HBSS+2% FBS vehicle were intraperitoneally (IP) injected using a 30 G insulin syringe (BD Biosciences cat# 328431). Transplanted cells were allowed 2 weeks to engraft prior to SCI.

### Generation of recombinant GST-Tcima

The *tcima* ORF was cloned into the BamHI/EcoRI sites in the pGEX-4T-1 backbone (Sigma cat# 28-9546-49). The zebrafish *tcima* ORF was PCR amplified from cDNA generated from 5 dpi wildtype SC using the following primers: *tcima*_ORF_F (gatc<u>GGATCC</u>ATGTCGAGCGACATGTACTC) and *tcima*_ORF_R (gatc<u>GAATTC</u> GCGCAGCTTCAGAAAGTTTC) where the underlined sequences indicate restriction enzyme sites. Once sequence confirmed, the resulting plasmid was transformed into BL21(DE3) bacteria. Bacterial cultures were grown until the OD600 reached 0.6-0.8. Expression was then induced with 1 mL of 100 mM IPTG for 5 hours. Pelleted bacteria were resuspended in PBS and sonicated under the following conditions: 15 seconds on, 45 seconds off at 45% input for a total of 6:15 minutes. The suspension was centrifuged for 30 minutes at 12,000 g at 4 °C. Glutathione Sepharose 4B beads (GE healthcare cat# 17-0756-01) were washed with 10 mL PBS then centrifuged at 1000 g at 4 °C for 1 minute for a total of three times. Supernatant was then incubated with pre-washed beads for 30 minutes at 4 °C. Beads were washed again with 10 mL PBS then centrifuged at 1000 g at 4 °C for 1 minute for 3 times. The slurry was then placed in a BioRad Poly-Prep Chromatography column (cat# 731-1550) and eluted in 2 mL of elution buffer (50 mM Tris-HCl pH 8.0, 10 mM glutathione). Dialysis was performed in dialysis tubing submersed in dialysis buffer (5 mM HEPES pH 7.6, 1 mM DTT, 0.2 mM PMSF, 1 mM EDTA, 10% glycerol) with gentle stirring at 4 °C overnight. Samples were aliquoted and stored at -80 °C.

### Immunoprecipitation and mass spectrometry

For protein lysates, 100 embryos at 2 dpf or 50 SCs at 3 dpi were used. Adult injured SCs were collected as described above. Samples were added to 4 °C lysis buffer (50 mM Tris-HCl pH 7.5, 125 mM NaCl, 1.5 mM MgCl, 1 mM EDTA, 5% glycerol, 0.4% NP-40, 0.1% Tween-20, 200 mM PMSF, and PI cocktail) and homogenized with a pestle. A second volume of lysis buffer was added and mixed by pipetting up and down. Samples were centrifuged at 16,000 g for 15 minutes at 4 °C. The supernatant was transferred to a new pre-chilled microfuge tube. 1/10 volume of NP-40 was added and the sample was mixed by vortexing. Samples were then centrifuged at 16,000 g for 15 minutes at 4 °C. The supernatant was transferred to a new pre-chilled microfuge tube and 4.25 µL of 100% glycerol was added per 10 µL of supernatant. Protein concentration was measured using Qubit and stored at -80 °C. The final volume of each lysate was 1 mL.

For immunoprecipitation, 50 µL (per sample) of Glutathione Sepharose 4B beads were washed with 10 mL PBS then centrifuged at 1000 g at 4 °C for 1 minute for a total of three times. GST-Tcima or GST alone were incubated with pre-washed GST-beads for 2 hours at 4 °C with gentle rocking. Samples were centrifuged at 1000 g for 3 minutes at 4 °C. The supernatant was removed, and beads were washed with 10 mL PBS+1% Triton X-100 then centrifuged at 1000 g at 4 °C for 1 minute for 3 times. The supernatant was removed and 100 µL of protein lysate was added. Samples were incubated 16 hours at 4 °C with gentle rocking. Samples were then centrifuged at 100 g for 3 minutes at 4 °C. Samples were washed three times for 5 minutes each at 4 °C with fresh wash buffer (5 mM HEPEs pH 7.5, 1 mM DTT, 0.2 mM PMSF, 1 mM EDTA, and 10% glycerol). Samples were resuspended in wash buffer and analyzed on the timsTOF Pro ion mobility mass spectrophotometer interfaced to a nanoElute LC system the Washington University Proteomics Shared Resource. Peptides with >5 spectral counts in the GST-Tcima samples and 0 spectral counts in GST-only samples were considered putative Tcima binding partners.

### Phagocytosis assays

For phagocytosis assays, 5 µL of 1 mg/mL Escherichia coli Bioparticles AlexaFluor-594 conjugate (ThermoFisher cat# E23370) were injected directly below the sealed wound at 3 dpi using a Hamilton syringe while fish were anesthetized. After 10 seconds, fish were returned to fresh fish water. Fish were collected at 16 hours after bioparticle injection for histology.

### Cell counting and statistical methods

All procedures and quantifications were performed blind to condition. For calculation of glial bridging, the cross-sectional Gfap^+^ area at the lesion core was normalized to the cross-sectional area of the intact spinal cord rostral to the lesion. For calculation of axon growth, biocytin quantification is described above. Cell counting was performed using a customized Fiji script (adapting ITCN: Image based Tool for counting nuclei, https://imagej.nih.gov/ij/plugins/itcn.html). This customized Fiji script incorporated user-defined inputs to define channels (including Hoechst) and to outline SC perimeters. To quantify nuclei, the following parameters were set in ITCN counter: width, 15; minimal distance, 7.5; threshold, 0.4. For each staining, thresholds were user-defined. Raw counts from Fiji were processed using a customized R Studio script. Absolute cell numbers are provided.

To quantify Mbp aggregation, a pixel intensity threshold value was determined for each experimental replicate based on an intensity threshold that included <1% of the whole imaged area in uninjured sections. This was to exclude quantification of myelinated axons, which had much lower signal intensity than Mbp aggregates that were present in injured fish. This threshold value was then used to identify the size of the imaged area that contained Mbp staining that was brighter than the set threshold. This quantified area was then labeled “Mbp^+^ Aggregate” area.

For volumetric quantifications of phagocytic cargo (both Mbp and bioparticles), Imaris was used. The Lcp1 channel was used to generate leukocyte surfaces. Any Lcp1^+^ surfaces <1000 pixels were removed to minimize background and bleed through of exceptionally strong Bioparticle puncta. The Lcp1^+^ surfaces were used to mask signal internal (engulfed) or external (extracellular) to leukocytes. Phagocytic cargo surfaces for each mask were then created using a single consistent threshold value across the entire experiment. Phagocytic cargo surfaces of <10 pixels were excluded to minimize background. The total volume of each mask was then calculated. The percent engulfment was calculated by dividing the internal engulfed volume by the total cargo (Mbp or bioparticles) volume for the image.

All sample sizes (n) are indicated for the number of animals used in each experiment, with specific sample sizes per each timepoint, genotype, and/or treatment indicated in figure legends. In cases where animals were lost due to death, the smallest sample size is indicated in the legend. In all graphs, dots indicate single animals and mean ± SEM are plotted. GraphPad Prism software was used to perform all statistical analysis. Unpaired student’s t-test (with Welch’s correction, whenever appropriate) or Mann Whitney test were used for comparing two groups. One-way or two-way ANOVA with appropriate corrections for multiple comparisons were used for comparing three or more groups. Specific statistical tests used are indicated in figure legends: p>0.05; *p<0.05, **p<0.01, and ***p< 0.001.

