## Supplementary material for "Elevated phagocytic capacity directs innate spinal cord repair": Supp Figures

### Figure S1

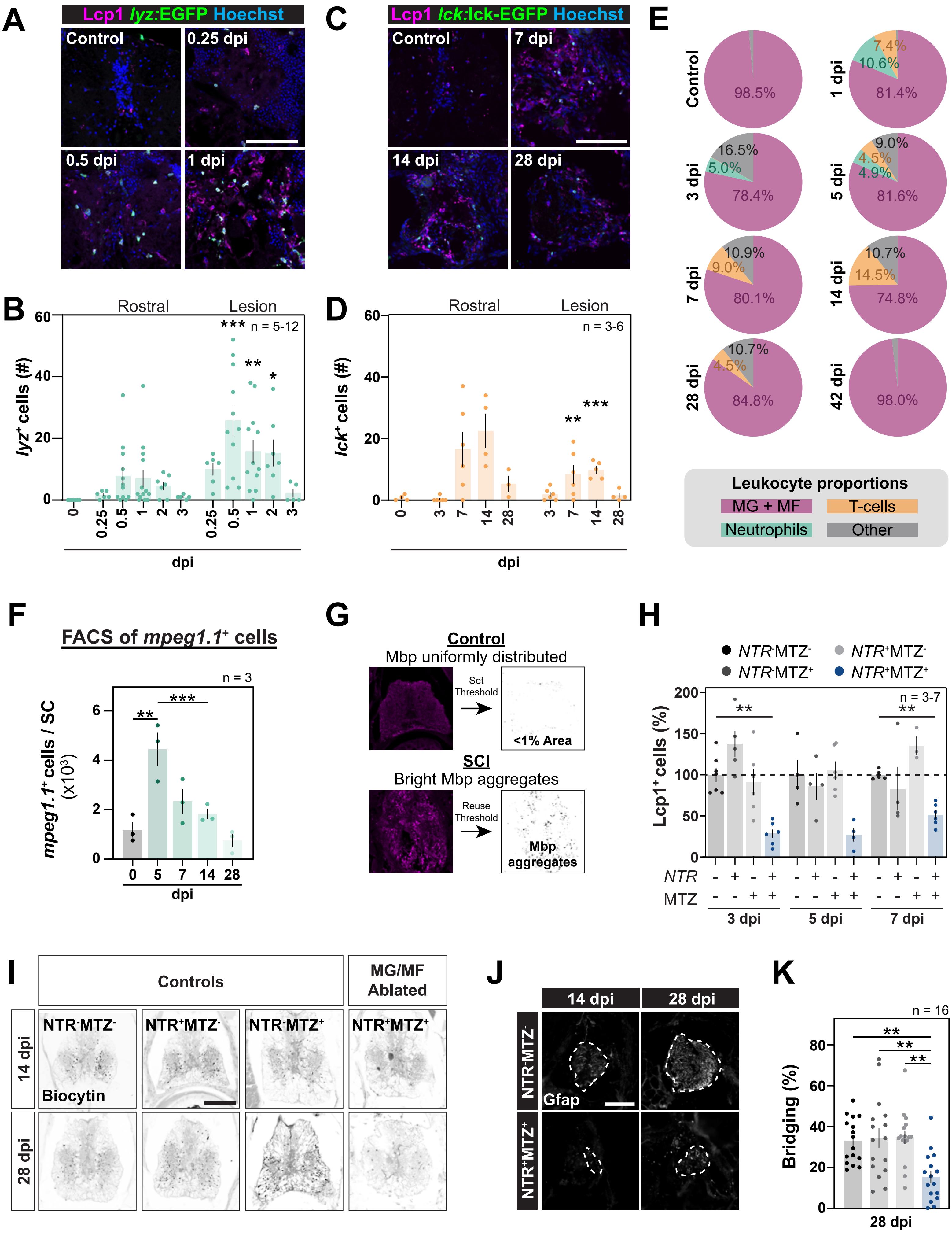
A
Lcp1 lyz:EGFP Hoechst
Control
0.25 dpi
0.5 dpi
1 dpi
B
Rostral
Lesion
n = 5-12
lyz+ cells (#)
dpi
C
Lcp1 lck:lck-EGFP Hoechst
Control
7 dpi
14 dpi
28 dpi
D
Rostral
Lesion
n = 3-6
lck+ cells (#)
dpi
E
Control
98.5%
1 dpi
7.4%
10.6%
81.4%
3 dpi
16.5%
5.0%
78.4%
5 dpi
9.0%
4.5%
4.9%
81.6%
7 dpi
10.9%
9.0%
80.1%
14 dpi
10.7%
14.5%
74.8%
28 dpi
10.7%
4.5%
84.8%
42 dpi
98.0%
Leukocyte proportions
MG + MF
T-cells
Neutrophils
Other
F
FACS of mpeg1.1+ cells
n = 3
mpeg1.1+ cells / SC (x10³)
dpi
G
Control
Mbp uniformly distributed
Set Threshold
<1% Area
SCI
Bright Mbp aggregates
Reuse Threshold
Mbp aggregates
H
NTR-MTZ-
NTR+MTZ-
NTR-MTZ+
NTR+MTZ+
n = 3-7
Lcp1+ cells (%)
NTR
MTZ
3 dpi
5 dpi
7 dpi
I
Controls
MG/MF Ablated
NTR-MTZ-
NTR+MTZ-
NTR-MTZ+
NTR+MTZ+
14 dpi
28 dpi
Biocytin
J
14 dpi
28 dpi
NTR-MTZ-
NTR+MTZ+
Gfap
K
n = 16
Bridging (%)
28 dpi

**Figure S1. Immune activation is necessary for SC regeneration (Related to Figure 1). (A,B)** Immunostaining for the pan-leukocyte marker L-plastin (Lcp1) in *Tg(lyz:EGFP)* zebrafish. Cross sections from uninjured controls or between 0.25 and 1 dpi are shown. Scale bar, 100 µm. Distance from the lesion is indicated in µm where negative and positive values indicate rostral and caudal levels, respectively. Sample sizes: Uninjured (6), 0.25 dpi (6), 0.5 dpi (11), 1 dpi (12), 2 dpi (7), 3 dpi (5). Two-way ANOVA (Dunnett's): *$p<0.05$, **$p<0.01$, ***$p<0.001$. Comparisons are to uninjured controls. **(C,D)** Immunostaining for the pan-leukocyte marker L-plastin (Lcp1) in *Tg(lck:lck-EGFP)* zebrafish. Cross sections from uninjured SCs (controls) or between 7 and 28 dpi are shown. Scale bar, 100 µm. Sample sizes: Uninjured (4), 3 dpi (6), 7 dpi (6), 14 dpi (4), 28 dpi (3). Two-way ANOVA (Dunnett's): **$p<0.01$, ***$p<0.001$. Comparisons are to uninjured control fish. **(E)** The proportion of $Lcp1^+$;*mpeg1.1:*$YFP^+$ leukocytes (microglia/macrophages, MGs + MF), *lyz*:$EGFP^+$ (neutrophils), *lck*:lck-$EGFP^+$ (pan-T-cells), or unlabeled (Other). **(F)** FACS of *Tg(mpeg1.1:YFP)* fish prior to SCI or at 5, 7, 14, and 28 dpi. The fraction of live cells that were *mpeg1.1*:$YFP^+$ are shown. Samples represent 4-5 SCs per replicate and 3 replicates were used for each time point. One-way ANOVA (Sidak's): **$p<0.01$, ***$p<0.001$. Comparisons are to uninjured controls, unless otherwise indicated. **(G)** Strategy to quantify Mbp aggregates following SCI. Mbp aggregates were labeled by defining a threshold for each experiment where the respective threshold labeled <1% of the total area in uninjured SC sections. This threshold was then used to count all pixels that were labeled with a higher intensity in injured sections. **(H)** To test the efficacy of MTZ treatment in *Tg(mpeg1.1:NTR-IRES-EGFP)* fish, animals were treated at 1 dpi with 5 mM MTZ followed by continued pulses of 1 mM MTZ every other day. The numbers of $Lcp1^+$ cells at 3, 5, and 7 dpi in MTZ- or vehicle-treated fish were normalized to time-matched transgene-negative vehicle-treated controls. Quantifications were performed at the lesion. Sample sizes: $NTR^-MTZ^-$ 3 dpi (7), $NTR^+MTZ^-$ 3 dpi (5), $NTR^-MTZ^+$ 3 dpi (6), $NTR^+MTZ^+$ 3 dpi (6), $NTR^-MTZ^-$ 5 dpi (4), $NTR^+MTZ^-$ 5 dpi (4), $NTR^-MTZ^+$ 5 dpi (6), $NTR^+MTZ^+$ 5 dpi (4), $NTR^-MTZ^-$ 7 dpi (5), $NTR^+MTZ^-$ 7 dpi (4), $NTR^-MTZ^+$ 7 dpi (3), $NTR^+MTZ^+$ 7 dpi (6). One-way ANOVA (Sidak's): **$p<0.01$. Comparisons are to time-matched $NTR^-MTZ^-$ controls. **(I)** Representative images from anterograde axon labeling of *Tg(mpeg1.1:NTR-IRES-EGFP)* MTZ- and vehicle-treated fish at 14 and 28 dpi. Rostral (-300 to -450 µm) images are shown. Scale bar, 100 µm. **(J,K)** Glial bridging at 14 or 28 dpi in microglia/macrophage-depleted fish. Representative images of the center of the $Gfap^+$ glial bridge in *Tg(mpeg1.1:NTR-IRES-EGFP)* MTZ- and vehicle-treated animals are shown. The cross-section of the glial bridge is outlined. Scale bar, 100 µm. Sample sizes: $NTR^-MTZ^-$ (16), $NTR^+MTZ^-$ (16), $NTR^-MTZ^+$ (16), $NTR^+MTZ^+$ (16). Two-way ANOVA (Dunnett's): **$p<0.01$. Comparisons are to $NTR^-MTZ^-$.

### Figure S2

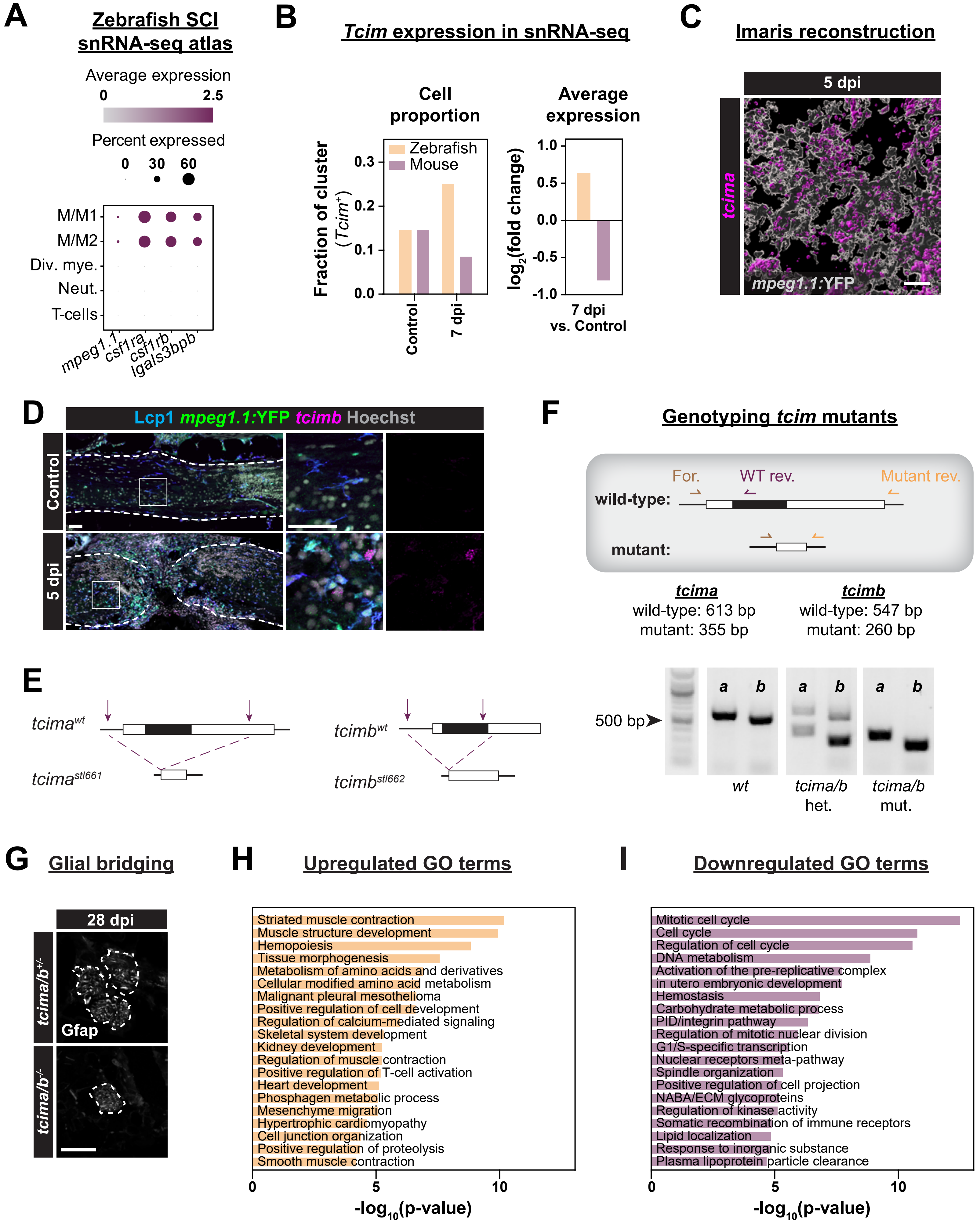

A
Zebrafish SCI snRNA-seq atlas
Average expression
0
2.5
Percent expressed
0 30 60
M/M1
M/M2
Div. mye.
Neut.
T-cells
mpeg1.1
csf1ra
csf1rb
lgals3bpb
B
Tcim expression in snRNA-seq
Cell proportion
Average expression
Fraction of cluster (Tcim+)
Zebrafish
Mouse
Control
7 dpi
log2(fold change)
7 dpi vs. Control
C
Imaris reconstruction
5 dpi
tcima
mpeg1.1:YFP
D
Lcp1 mpeg1.1:YFP tcimb Hoechst
Control
5 dpi
F
Genotyping tcim mutants
For.
WT rev.
Mutant rev.
wild-type:
mutant:
tcima
wild-type: 613 bp
mutant: 355 bp
tcimb
wild-type: 547 bp
mutant: 260 bp
500 bp
a b a b a b
wt
tcima/b het.
tcima/b mut.
E
tcima^wt
tcima^stl661
tcimb^wt
tcimb^stl662
G
Glial bridging
28 dpi
tcima/b+/-
tcima/b-/-
Gfap
H
Upregulated GO terms
Striated muscle contraction
Muscle structure development
Hemopoiesis
Tissue morphogenesis
Metabolism of amino acids and derivatives
Cellular modified amino acid metabolism
Malignant pleural mesothelioma
Positive regulation of cell development
Regulation of calcium-mediated signaling
Skeletal system development
Kidney development
Regulation of muscle contraction
Positive regulation of T-cell activation
Heart development
Phosphagen metabolic process
Mesenchyme migration
Hypertrophic cardiomyopathy
Cell junction organization
Positive regulation of proteolysis
Smooth muscle contraction
-log10(p-value)
I
Downregulated GO terms
Mitotic cell cycle
Cell cycle
Regulation of cell cycle
DNA metabolism
Activation of the pre-replicative complex
in utero embryonic development
Hemostasis
Carbohydrate metabolic process
PID/integrin pathway
Regulation of mitotic nuclear division
G1/S-specific transcription
Nuclear receptors meta-pathway
Spindle organization
Positive regulation of cell projection
NABA/ECM glycoproteins
Regulation of kinase activity
Somatic recombination of immune receptors
Lipid localization
Response to inorganic substance
Plasma lipoprotein particle clearance
-log10(p-value)

**Figure S2. *tcima/b* mutants display poor metrics of SC regeneration (Related to Figure 2). (A)** Dot plots from snRNA-seq zebrafish SCI atlas (Saraswathy *et al.*, 2023). The microglia/macrophage cluster 1 (M/M1) was used to identify top microglia/macrophage marker genes because this cluster was the largest microglia/macrophage cluster that expressed the canonical marker genes *mpeg1.1, csf1ra, csf1rb,* and *lgals3bpb*. **(B)** Expression of zebrafish *tcima* (gold) or mouse *Tcim* (magenta) in snRNA-seq or scRNA-seq datasets. **(C)** Imaris 3D reconstructions of *tcima* expression in *mpeg1.1:*YFP$^{+}$ cells. Scale bar, 10 µm. **(D)** HCR *in situ* hybridization of *tcimb* and Lcp1 staining were performed on *Tg(mpeg1.1:YFP)* fish. Sagittal SC sections from uninjured or 5 dpi animals were used. Scale bars, 50 µm. **(E)** CRISPR/Cas9 was used to excise the entire coding sequence of either *tcima* or *tcimb* to generate loss-of-function mutants. CRISPR/Cas9 guide RNA target sites (purple arrows), exons (black boxes); untranslated regions (UTRs, white boxes). **(F)** Genotyping schematic and representative results for *tcima* (*a*) and *tcimb* (*b*) deletions. For each gene, three primers were designed such that PCR would produce one larger band for wild-type product (613 bp for *tcima* and 547 bp for *tcimb*) and one smaller band for mutant product (355 bp for *tcima* and 260 bp for *tcimb*). The 100 bp DNA ladder was used, 500 bp reference band indicated. **(G)** Representative images of the center of the Gfap$^{+}$ glial bridge in control siblings and *tcima/b* mutants at 28 dpi. The cross-section of the glial bridge is outlined. Scale bar, 100 µm. **(H,I)** Gene ontology (GO) analysis for all statistically significant genes identified upregulated (H) or downregulated (I) in *tcima/b* mutants compared to control siblings.

### Figure S3

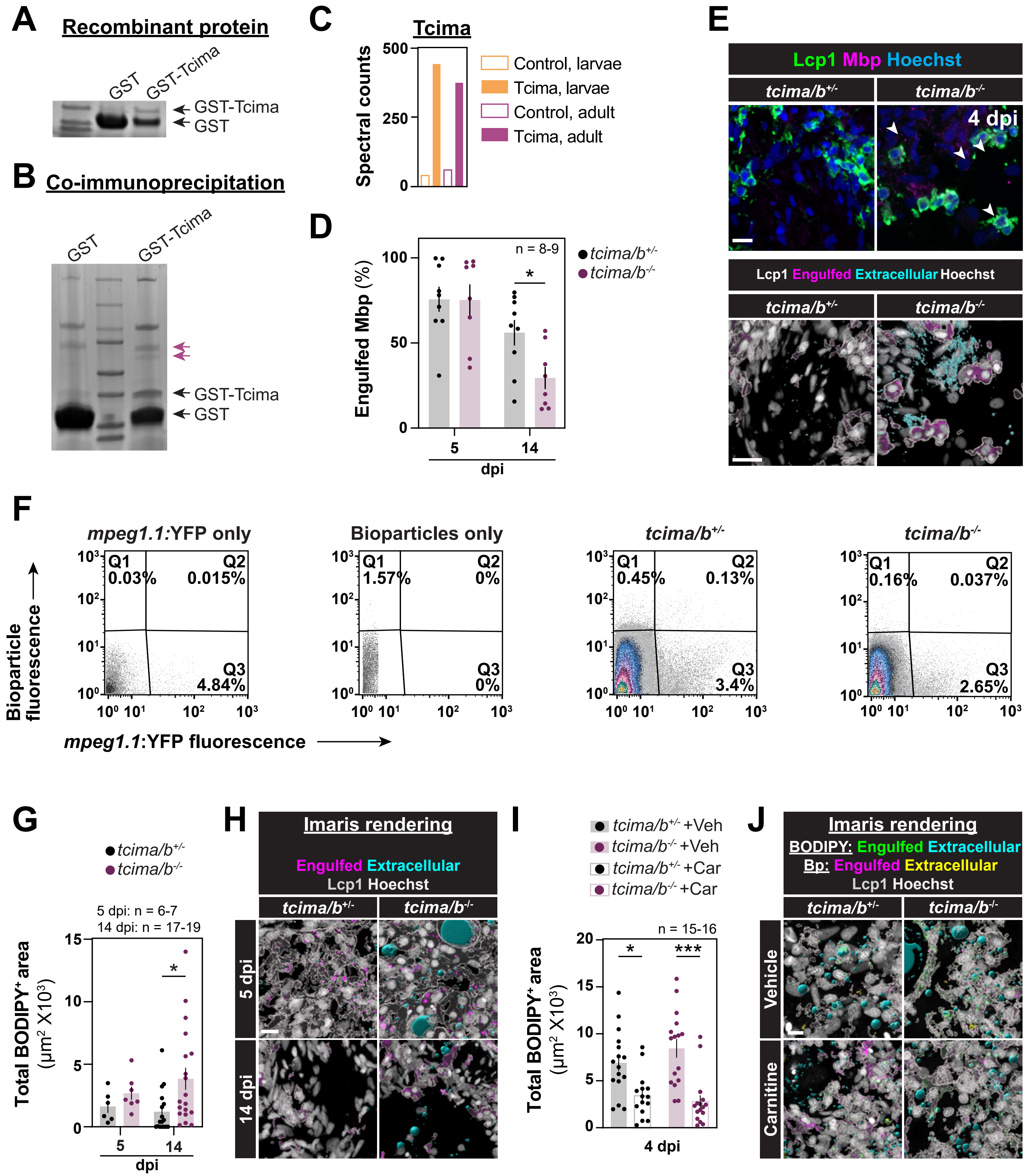

A
Recombinant protein
GST
GST-Tcima
GST-Tcima
GST
B
Co-immunoprecipitation
GST
GST-Tcima
GST-Tcima
GST
C
Tcima
Spectral counts
500
250
0
Control, larvae
Tcima, larvae
Control, adult
Tcima, adult
D
Engulfed Mbp (%)
100
50
0
n = 8-9
*
5
14
dpi
tcima/b+/-
tcima/b-/-
E
Lcp1 Mbp Hoechst
tcima/b+/-
tcima/b-/-
4 dpi
Lcp1 Engulfed Extracellular Hoechst
tcima/b+/-
tcima/b-/-
F
mpeg1.1:YFP only
Bioparticles only
tcima/b+/-
tcima/b-/-
Q1 0.03%
Q2 0.015%
Q3 4.84%
Q1 1.57%
Q2 0%
Q3 0%
Q1 0.45%
Q2 0.13%
Q3 3.4%
Q1 0.16%
Q2 0.037%
Q3 2.65%
Bioparticle fluorescence
mpeg1.1:YFP fluorescence
G
tcima/b+/-
tcima/b-/-
5 dpi: n = 6-7
14 dpi: n = 17-19
Total BODIPY+ area (μm2 X103)
15
10
5
0
*
5
14
dpi
H
Imaris rendering
Engulfed Extracellular
Lcp1 Hoechst
tcima/b+/-
tcima/b-/-
5 dpi
14 dpi
I
tcima/b+/- +Veh
tcima/b-/- +Veh
tcima/b+/- +Car
tcima/b-/- +Car
n = 15-16
Total BODIPY+ area (μm2 X103)
20
15
10
5
0
*
***
4 dpi
J
Imaris rendering
BODIPY: Engulfed Extracellular
Bp: Engulfed Extracellular
Lcp1 Hoechst
tcima/b+/-
tcima/b-/-
Vehicle
Carnitine

**Figure S3. Tcim is necessary for debris clearance following SCI (Related to Figure 4). (A,B)** Recombinant GST-Tcima was generated (A) and used to perform immunoprecipitation (Co-IP, B) with protein lysate collected from whole 2 dpf larvae or 3 dpi adult SC. Purified products from Co-IP were ran on a polyacrylamide gel and stained with Coomassie Blue. The expected sizes of GST and GST-Tcima are indicated. Additional bands present after Co-IP with GST-Tcima (magenta arrows). **(C)** Spectral counts of Tcima in GST and GST-Tcima samples. **(D,E)** Mbp and Lcp1 staining in *tcima/b*$^{-/-}$ and control fish. Cross sections from 5 and 14 dpi are shown. Imaris was used to partition bioparticles into Extracellular (cyan) or Engulfed (magenta) channels based on Lcp1 surfaces (gray). Scale bars, 10 µm. Sample sizes: *tcima/b*$^{+/-}$ 5 dpi (9), *tcima/b*$^{-/-}$ 5 dpi (8), *tcima/b*$^{+/-}$ 14 dpi (9), *tcima/b*$^{-/-}$ 14 dpi (8). One-way ANOVA (Sidak's): *p<0.05. **(F)** FACS of *Tg(mpeg1.1:YFP); tcima/b*$^{-/-}$ and *Tg(mpeg1.1:YFP); tcima/b*$^{+/-}$ fish that were injected with AlexaFluor-conjugated E. coli bioparticles at 3 dpi. Samples were collected 16 hours after bioparticle injection. FACS plots shown are from one representative replicate for each genotype and the controls used to set fluorescence gates (*mpeg1.1*:YFP only and bioparticles only). **(G,H)** Imaris reconstructions and quantification of BODIPY and Lcp1 staining for lipids and pan-leukocytes, respectively, in *tcima/b* mutants and control siblings at 5 and 14 dpi. Scale bar, 10 µm. Sample sizes: *tcima/b*$^{+/-}$ 5 dpi (6), *tcima/b*$^{-/-}$ 5 dpi (7), *tcima/b*$^{+/-}$ 14 dpi (17), *tcima/b*$^{-/-}$ 14 dpi (19). One-way ANOVA (Sidak's): *p<0.05. **(I,J)** Quantification for histology of carnitine- or vehicle-injected *tcima/b* mutants and control siblings stained with BODIPY and Lcp1. Animals were also injected at 3 dpi with fluorescently-labeled bioparticles, and collected 16 hours after injection for histology. Scale bar, 10 µm. Sample sizes: *tcima/b*$^{+/-}$ vehicle (16), *tcima/b*$^{+/-}$ L-carnitine (15), *tcima/b*$^{-/-}$ vehicle (16), *tcima/b*$^{-/-}$ L-carnitine (16). Two-way ANOVA (Tukey's): *p<0.05, ***p<0.001.

### Figure S4

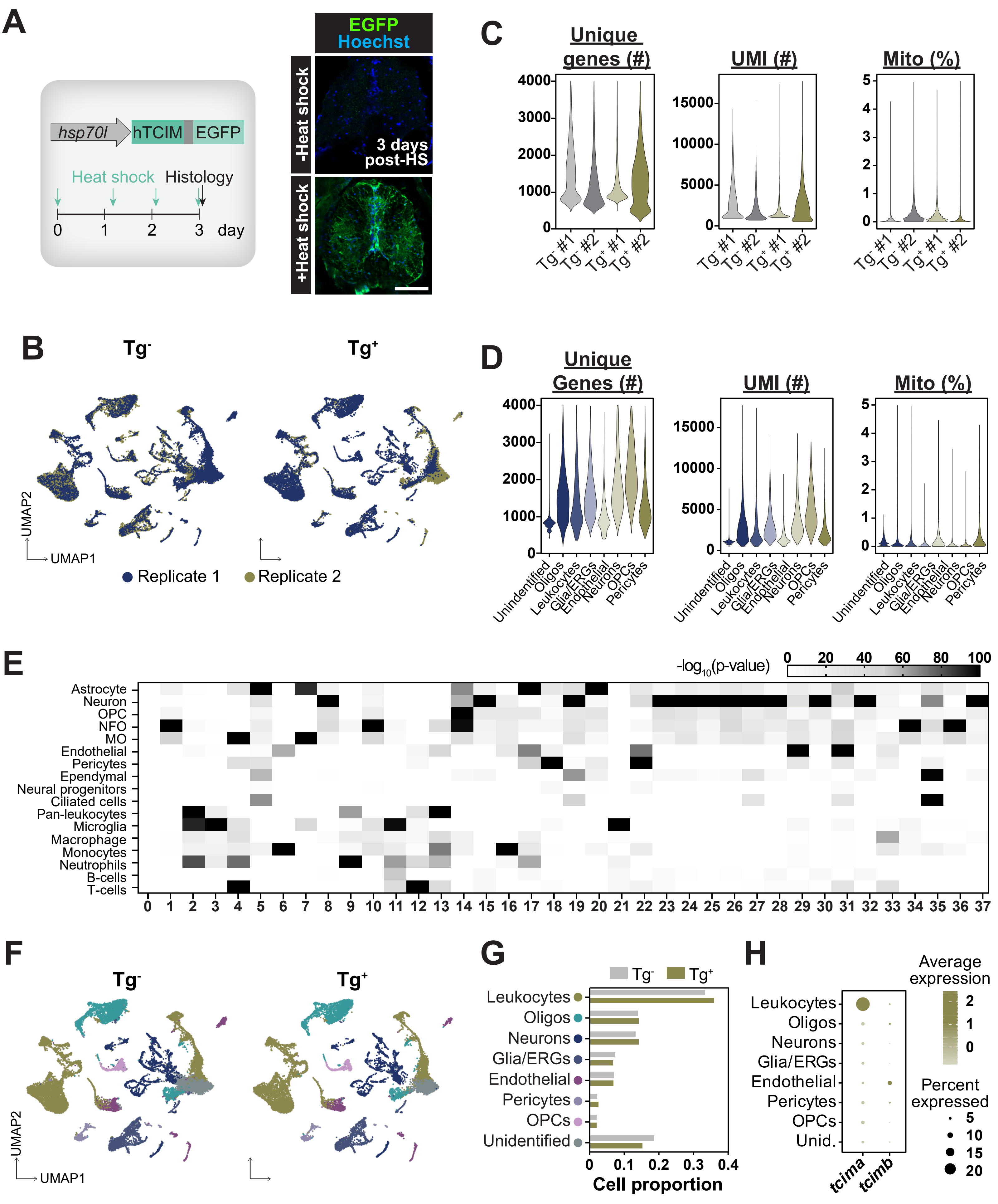

A
hsp70l
hTCIM
EGFP
Heat shock
Histology
0
1
2
3
day
EGFP
Hoechst
-Heat shock
+Heat shock
3 days post-HS
B
Tg-
Tg+
UMAP2
UMAP1
Replicate 1
Replicate 2
C
Unique genes (#)
UMI (#)
Mito (%)
Tg- #1
Tg- #2
Tg+ #1
Tg+ #2
D
Unique Genes (#)
UMI (#)
Mito (%)
Unidentified
Oligos
Leukocytes
Glia/ERGs
Endothelial
Neurons
OPCs
Pericytes
E
-log10(p-value)
Astrocyte
Neuron
OPC
NFO
MO
Endothelial
Pericytes
Ependymal
Neural progenitors
Ciliated cells
Pan-leukocytes
Microglia
Macrophage
Monocytes
Neutrophils
B-cells
T-cells
F
Tg-
Tg+
UMAP2
UMAP1
G
Tg-
Tg+
Leukocytes
Oligos
Neurons
Glia/ERGs
Endothelial
Pericytes
OPCs
Unidentified
Cell proportion
H
Leukocytes
Oligos
Neurons
Glia/ERGs
Endothelial
Pericytes
OPCs
Unid.
tcima
tcimb
Average expression
Percent expressed

**Figure S4. TCIM expression requires microglia/macrophages and does not alter cell composition at the pan-neural level (Related to Figure 6). (A)** *Tg(hsp70l:hTCIM-2A-EGFP)* uninjured fish treated with or without three days of daily heat shock and stained for EGFP. Scale bar, 100 µm. **(B)** UMAPs of biological replicates for TCIM-expressing and control siblings. **(C,D)** Quality control metrics for snRNA-seq in TCIM-expressing and control transgene-negative siblings. The numbers of unique genes, unique molecular identifiers (UMI), and percentage of mitochondrial genes (Mito) are shown. Each sample count matrix was filtered for genes that were expressed in at least three cells, for the number of unique genes per cell (expressing 200 to 4000 unique genes), and for mitochondrial gene percentage (<5). Sequencing was performed in biological duplicates. Each duplicate represents 35-39 SCs. **(E)** Cell subpopulations were identified using hypergeometric scoring against the VNM database, (Saraswathy *et al.*, 2023). Normalized heatmap of overlap between subcluster top marker genes and the VNM database is shown. Oligodendrocyte precursor cell (OPC), newly formed oligodendrocyte (NFO), myelinating oligodendrocyte (MO). **(F)** UMAP representation of neural subclusters in TCIM-expressing and control fish. **(G)** Cell proportions based on subclustering. Oligodendrocytes (Oligos), ependymal radial glia (ERGs), oligodendrocyte precursor cell (OPCs). **(H)** Dot plot showing expression of *tcima* and *tcimb* in neural cells.

### Figure S5

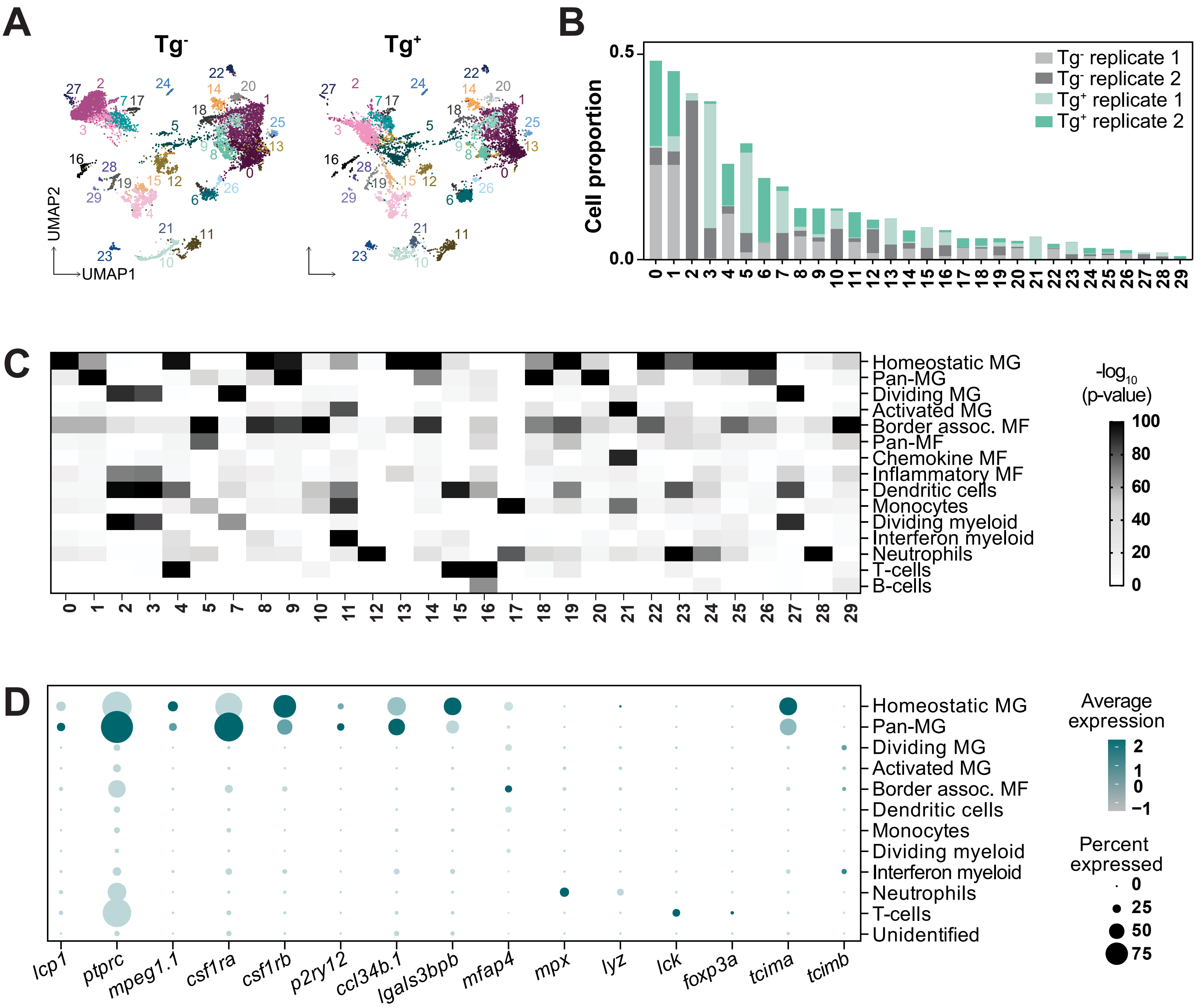
A
Tg-
Tg+
UMAP2
UMAP1
B
Cell proportion
Tg- replicate 1
Tg- replicate 2
Tg+ replicate 1
Tg+ replicate 2
C
Homeostatic MG
Pan-MG
Dividing MG
Activated MG
Border assoc. MF
Pan-MF
Chemokine MF
Inflammatory MF
Dendritic cells
Monocytes
Dividing myeloid
Interferon myeloid
Neutrophils
T-cells
B-cells
-log10 (p-value)
D
Unidentified
Average expression
Percent expressed
lcp1
ptprc
mpeg1.1
csf1ra
csf1rb
p2ry12
ccl34b.1
lgals3bpb
mfap4
mpx
lyz
lck
foxp3a
tcima
tcimb

**Figure S5. TCIM expression reprograms myeloid precursors into mature phagocytes (Related to Figure 6). (A)** UMAPs of leukocyte subclusters. A resolution of 0.7 was selected as this was the resolution where subclusters became stable. **(B)** Leukocyte subcluster proportions across replicates in TCIM-expressing and control SCs. Leukocyte nuclei were separated into 30 subclusters. **(C)** Leukocyte cell types from the VNM database were used to identify the 30 leukocyte subclusters. Normalized heatmap of overlap between leukocyte subcluster top marker genes and the leukocyte atlas is shown. Genes used for the leukocyte atlas can be found in Table S8. **(D)** Dot plots for known markers of leukocyte subtypes, including pan-leukocytes (*lcp1*), myeloid cells (*ptprc*), microglia/macrophages (*mpeg1.1, csf1ra, and csf1rb*), microglia (*p2ry12, ccl34b.1, and lgals3bpb*), macrophages (*mfap4*), neutrophils (*mpx and lyz*), and T-cells (*lck and foxp3a*). Expression of *tcima* and *tcimb* are also shown.

### Figure S6

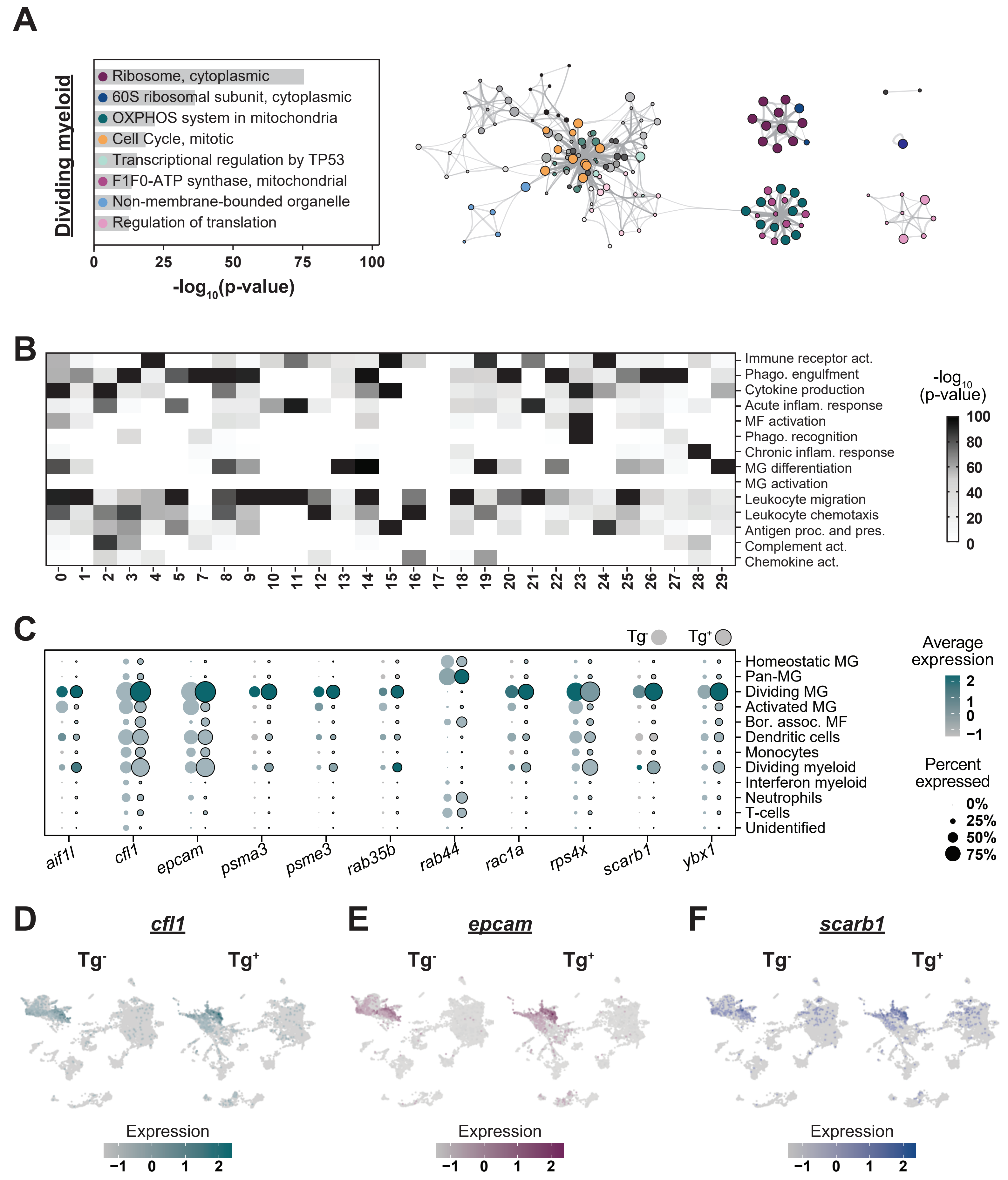
A
Dividing myeloid
Ribosome, cytoplasmic
60S ribosomal subunit, cytoplasmic
OXPHOS system in mitochondria
Cell Cycle, mitotic
Transcriptional regulation by TP53
F1F0-ATP synthase, mitochondrial
Non-membrane-bounded organelle
Regulation of translation
0 25 50 75 100
-log10(p-value)
B
Immune receptor act.
Phago. engulfment
Cytokine production
Acute inflam. response
MF activation
Phago. recognition
Chronic inflam. response
MG differentiation
MG activation
Leukocyte migration
Leukocyte chemotaxis
Antigen proc. and pres.
Complement act.
Chemokine act.
0 1 2 3 4 5 7 8 9 10 11 12 13 14 15 16 17 18 19 20 21 22 23 24 25 26 27 28 29
-log10 (p-value)
100 80 60 40 20 0
C
Tg- Tg+
Homeostatic MG
Pan-MG
Dividing MG
Activated MG
Bor. assoc. MF
Dendritic cells
Monocytes
Dividing myeloid
Interferon myeloid
Neutrophils
T-cells
Unidentified
aif1l cfl1 epcam psma3 psme3 rab35b rab44 rac1a rps4x scarb1 ybx1
Average expression
2 1 0 −1
Percent expressed
0% 25% 50% 75%
D
cfl1
Tg- Tg+
Expression
−1 0 1 2
E
epcam
Tg- Tg+
Expression
−1 0 1 2
F
scarb1
Tg- Tg+
Expression
−1 0 1 2

**Figure S6. Leukocytes in TCIM-expressing fish are enriched for phagocytic genes (Related to Figure 6). (A)** GO analysis was performed on top marker genes identified in the dividing myeloid subclusters. Genes that were found as a top marker were used to produce GO term lists from dividing myeloid subclusters. Metascape GO term clustering schematics are shown. Node size is proportional to the number of input genes falling into each GO term, and the line weight indicates the similarity in gene lists between connected GO terms. **(B)** A hypergeometric probability test was used to identify the most statistically significant GO term associated with each leukocyte subcluster by analyzing the overlap between subcluster top marker genes and the immune GO term database. Gene lists used for analysis are in Table S12. To generate the Immune GO Term database, we utilized the gene ontology database Amigo. Inclusion criteria for GO terms is described in the methods section. Top marker genes from each subcluster were compared to the Immune GO Term database to generate a normalized heatmap. **(C)** Dot plots for genes that fell within the "phagocytosis engulfment" GO term. **(D-F)** Feature plots of *cfl1, epcam*, and *scarb1* for control or TCIM-expressing samples.
